# Co-obligate symbioses have repeatedly evolved across aphids, but partner identity and nutritional contributions vary across lineages

**DOI:** 10.1101/2022.08.28.505559

**Authors:** Alejandro Manzano-Marín, Armelle Coeur d’acier, Anne-Laure Clamens, Corinne Cruaud, Valérie Barbe, Emmanuelle Jousselin

## Abstract

Aphids are a large family of phloem-sap feeders. They typically rely on a single bacterial endosymbiont, *Buchnera aphidicola*, to supply them with essential nutrients lacking in their diet. This association with *Buchnera* was described in model aphid species from the Aphidinae subfamily and has been assumed to be representative of most aphids. However, in two lineages, *Buchnera* has lost some essential symbiotic functions and is now complemented by additional symbionts. Though these cases break our view of aphids harbouring a single obligate endosymbiont, we know little about the extent, nature, and evolution of these associations across aphid subfamilies. Here, using metagenomics on 25 aphid species from nine subfamilies, re-assembly and re-annotation of 20 aphid symbionts previously sequenced, and 16S rRNA amplicon sequencing on 223 aphid samples (147 species from 12 subfamilies), we show that dual symbioses have evolved anew at least six times. We also show that these secondary co-obligate symbionts have typically evolved from facultative symbiotic taxa. Genome-based metabolic inference confirms interdependencies between *Buchnera* and its partners for the production of essential nutrients but shows contributions vary across pairs of co-obligate associates. Fluorescent *in situ* hybridisation microscopy shows a common bacteriocyte localisation of two newly acquired symbionts. Lastly, patterns of *Buchnera* genome evolution reveal that small losses affecting a few key genes can be the onset of these dual systems, while large gene losses can occur without any co-obligate symbiont acquisition. Hence, the *Buchnera*-aphid association, often thought of as exclusive, seems more flexible, with a few metabolic losses having recurrently promoted the establishment of a new co-obligate symbiotic partner.

## Introduction

Nutritional symbioses between animals and microorganisms are widespread in nature, and are particularly present in hemipteran insects with a nutrient-restricted diet. Aphids (Hemiptera: Apdididae) are a family of around 5,200 species organised into 23 extant subfamilies (Favret, 2022). Their diet consists entirely of plant phloem, which is rich in sugars, but poor in essential amino acids and B vitamins (Douglas, 2006; Sandstrom and Moran, 1999; Ziegler, 1975). In order to overcome this limitation, aphids are associated to an obligate vertically-transmitted endosymbiotic bacteria, *Buchnera aphidicola*, that synthesises nutrients lacking in their diet, namely essential amino acids (EAAs) and B vitamins (Baumann, 2005; Bermingham *et al*., 2009; Blow *et al*., 2020; Douglas, 1998; Hansen and Moran, 2011; Shigenobu *et al*., 2000). The long-term association of *Buchnera* and aphids is evidenced by the high degree of *Buchnera* genome synteny and general congruence of aphid and symbiont phylogenies (Baumann *et al*., 1995; Funk *et al*., 2000; Jousselin *et al*., 2009; Moran *et al*., 1993; Nováková *et al*., 2013; van Ham *et al*., 2003). As a result of this ancient association, *Buchnera* strains have evolved highly reduced, AT-rich, and gene-dense genomes (Chong and Moran, 2018). While genome erosion has affected many functional categories, extant *Buchnera* strains have markedly retained genes involved in the biosynthesis of essential amino acids and B vitamins. However intimate the aphid-*Buchnera* symbiosis is, long-lived associations can break down, leading to symbiont replacement or complementation.

While ancient symbiont replacement and complementation has been well documented in hemipteran taxa such as the Auchenorrhyncha (*e.g.* cicadas, treehoppers, leafhoppers, and planthoppers; Koga and Moran, 2014; Łukasik *et al*., 2018; Matsuura *et al*., 2018; McCutcheon and Moran, 2007; Michalik *et al*., 2021), Pseudococcinae (mealybugs; Husnik and McCutcheon, 2016; Szabó *et al*., 2017), Adelgidae (adelgids; Dial *et al*., 2022; Szabó *et al*., 2022; Toenshoff *et al*., 2012; Weglarz *et al*., 2018), and Psylloidea (psyllids; Nakabachi *et al*., 2013, 2020; Sloan and Moran, 2012), these associations have not been widely reported across aphids. The mutualism between aphids and *Buchnera* has generally been seen as stable and quite exclusive. However, most of our knowledge on this symbiosis comes from one aphid lineage: the Aphidinae subfamily. This subfamily encompasses about 3,150 species (around 60% of aphid diversity) including most aphid pests, and thus, has been the one that has been most studied. Yet, microbial metagenomic data from aphids outside Aphidinae is revealing that the aphid/*Buchnera* relationship might not be as stable as widely thought. In *Geopemphigus* and within Cerataphidini aphids, *Buchnera* has been lost, and in its place, *Bacteroidota* (Chong and Moran, 2018) and yeast-like symbionts are found in these species (Fukatsu and Ishikawa, 1992; Fukatsu *et al*., 1994), respectively. In both cases, the new symbionts have the genetic capacity to take over the nutrient provisioning role of the now defunct *Buchnera* (Chong and Moran, 2018; Vogel and Moran, 2013), which would allow the aphids to continue thriving on a nutrient-deficient diet. In addition to symbiont replacement, symbiont complementation can arise if a co-existing microbe has the metabolic capacity to rescue or take over one or more of the roles of the pre-existing associate. Such co-obligate symbioses have indeed arose in at least two aphid groups: the Lachninae (Manzano-Marín *et al*., 2017) and the *Periphyllus* genus (Chaitophorinae: Chaitophorini; Monnin *et al*., 2020; Renoz *et al*., 2022a). Very recently, an additional two reports of co-obligate symbioses have been published in two species from two different tribes: *Sipha maydis* (Chaitophorinae: Siphini; Renoz *et al*., 2022b), which has been idenpendently analysed in the current work, and *Ceratovacuna japonica* (Hormaphidinae: Cerataphidini; Yorimoto *et al*., 2022). These secondary co-obligate symbionts now complement evolved auxotrophies of their corresponding *Buchnera* partners, most commonly those for biotin and riboflavin and less often those for tryptophan and histidine. While Monnin *et al*. (2020) also reported alleged co-obligate symbiotic systems in two Aphidinae species, post-publication re-analysis of their work has cast important doubts regarding some of their analyses and interpretation of results (Manzano-Marín, 2020). Thus, these co-obligate associations within Aphidinae are not further discussed in the context of the current work.

In the remaining aphid subfamilies, there is no genomic evidence for the occurrence of multi-partner symbioses. However, a series of microscopic studies have revealed co-occurring bacteriocyte-associated bacteria outside of the Lachninae subfamily and *Periphyllus* genus which show co-obligate like characteristics. Early microscopic evidence showed co-existing bacteria in separate bacteriocytes to those of *Buchnera* in the aphids *Drepanosiphum* sp. (Drepanosiphinae), *Periphyllus testudinaceus* (Chaitophorinae: Chaitophorini), and *Panaphis juglandis* (Calaphidinae: Panaphidini) (Buchner, 1953). More recent work has also shown co-obligate like organisms in *Yamatocallis* spp. (Drepanosiphinae) (Fukatsu, 2001; Fukatsu and Ishikawa, 1993), *Sipha maydis* (Chaitophorinae: Siphini), *Anoecia corni* (Anoeciinae), and *Glyphina betulae* (Thelaxinae) (Kot, 2012; Michalik, 2010; Michalik *et al*., 2014). In most above-mentioned cases, the co-obligate symbionts are not only inhabiting their own bacteriocytes, but also show a spherical cell shape, which is characteristic of many obligate symbionts of aphids and adelgids with drastically reduced genomes (Dial *et al*., 2022; Lamelas *et al*., 2008, 2011b; Manzano-Marín *et al*., 2016, 2017, 2020; Michalik *et al*., 2021; Szabó *et al*., 2022; Toenshoff *et al*., 2012, 2014).

In this work we sought to explore the extent, nature, identity, and metabolic capabilities of nutritional endosymbiotic consortia across aphids. For this purpose, we assembled the most comprehensive and diverse set of aphid symbionts to date. This dataset included 25 newly sequenced symbiont genomes as well as 20 re-assembled and/or re-annotated previously sequenced ones. Through high-throughput 16S rRNA gene amplicon sequencing of 147 species of aphids, we were able to corroborate the identity of these symbionts, their prevalence across species from the same lineages, and explored whether new obligate symbionts necessarily emerge from frequent symbionts in the microbiota. In addition, through genome-based metabolic inference we explored the co-dependency of the co-existing symbionts and collaboration for the production of their hosts’ essential nutrients. Through phylogenetic analyses we investigated the origin of co-obligate symbiont lineages. Lastly, using of Fluorescence *in situ* hybridisation (FISH), we corroborate the identity and bacteriocyte-specific localisation of two distantly-related secondary co-obligate symbionts in selected aphid species.

## Results

### *Buchnera* genome has repeatedly undergone further genome reduction

In order to reconstruct a comprehensive phylogeny of *Buchnera aphidicola* (hereafter *Buchnera*) to aid in our evolutionary interpretations, we assembled a genomic dataset of 48 strains housed by different aphid species (supplementary table S1, Supplementary Material online). This dataset represents aphid symbionts from 13 different subfamilies, including the most speciose ones, and is, to our knowledge, the largest and most diverse dataset assembled for *Buchnera*. As noted previously by Chong *et al*. (2019), *Buchnera* from Aphidinae have among the largest genome sizes as well as highest G+C content and number of coding sequences (CDSs, figure 1). Conversely, the genomes of Lachninae, known to host co-obligate symbionts (Manzano-Marín *et al*., 2017, 2020; Meseguer *et al*., 2017), have among the smallest genomes with under 400 CDSs and lower G+C contents. With our expanded dataset, we have also observed many genomes have intermediate values between those of Aphidinae and Lachninae, namely those of Chaitophorinae, Thelaxinae, Neophyllaphidinae, Anoeciinae, Mindarinae, and one Eriosomatini.

**Figure 1.**
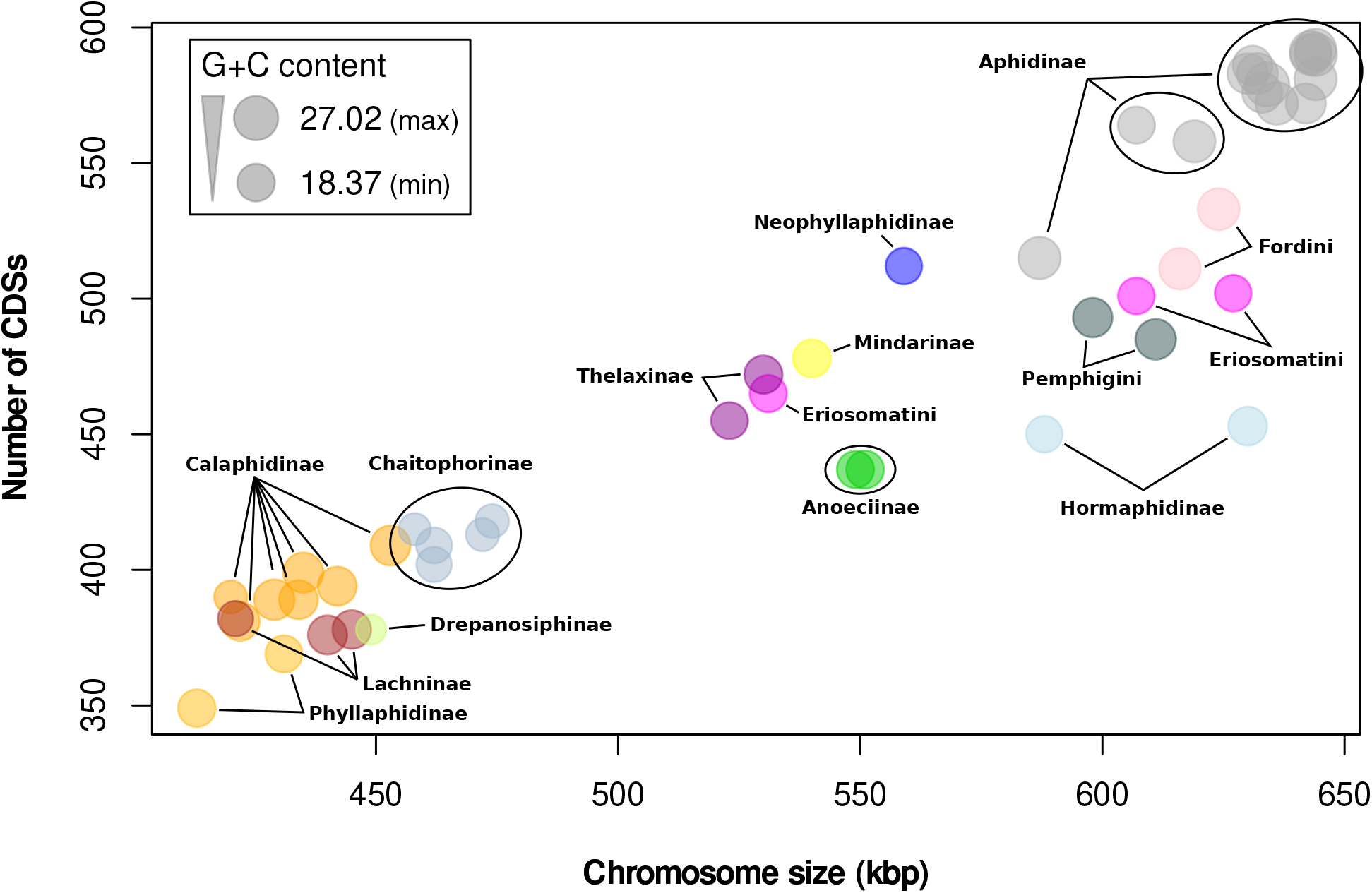
Relationship of genome size *vs.* CDS content in *Buchnera* genomes. Scatter plot illustrating the diversity of genome size, number of CDSs and G+C content of selected *Buchnera* genomes. Taxonomic classification of aphid host is displayed in bold black lettering. Dots are colour coded according to the host’s subfamily or tribal classification. Size of circles represent the G+C content as shown in the legend box scale key.

After thorough manual curation of the *Buchnera* genomes, we extracted the 229 single-copy core proteins and used them to reconstruct a concatenated protein phylogeny for *Buchnera* (figure 2). As in previous works (Chen *et al*., 2017; Li *et al*., 2014; Nováková et al., 2013; Ortiz-Rivas and Martínez-Torres, 2010), we found support for the non-monophyly of the Eriosomatinae subfamily. Also, the symbionts of both sequenced Phyllaphidinae species were recovered nested within those of Calaphidinae, which corroborates previous results recovering members of this subfamily as closely related to species or tribes of Calaphidinae (Nováková et al., 2013; von Dohlen and Moran, 2000). It is important to note that there remains some uncertainty regarding the position of the root, given the long branch leading to the *Buchnera* lineage. However, alternative phylogenetic reconstructions with and without the more distantly related *Escherichia coli* as an outgroup, gave identical topologies with small variations in support for some internal branching (alternative trees found in https://doi.org/10.5281/zenodo.6394197).

**Figure 2.**
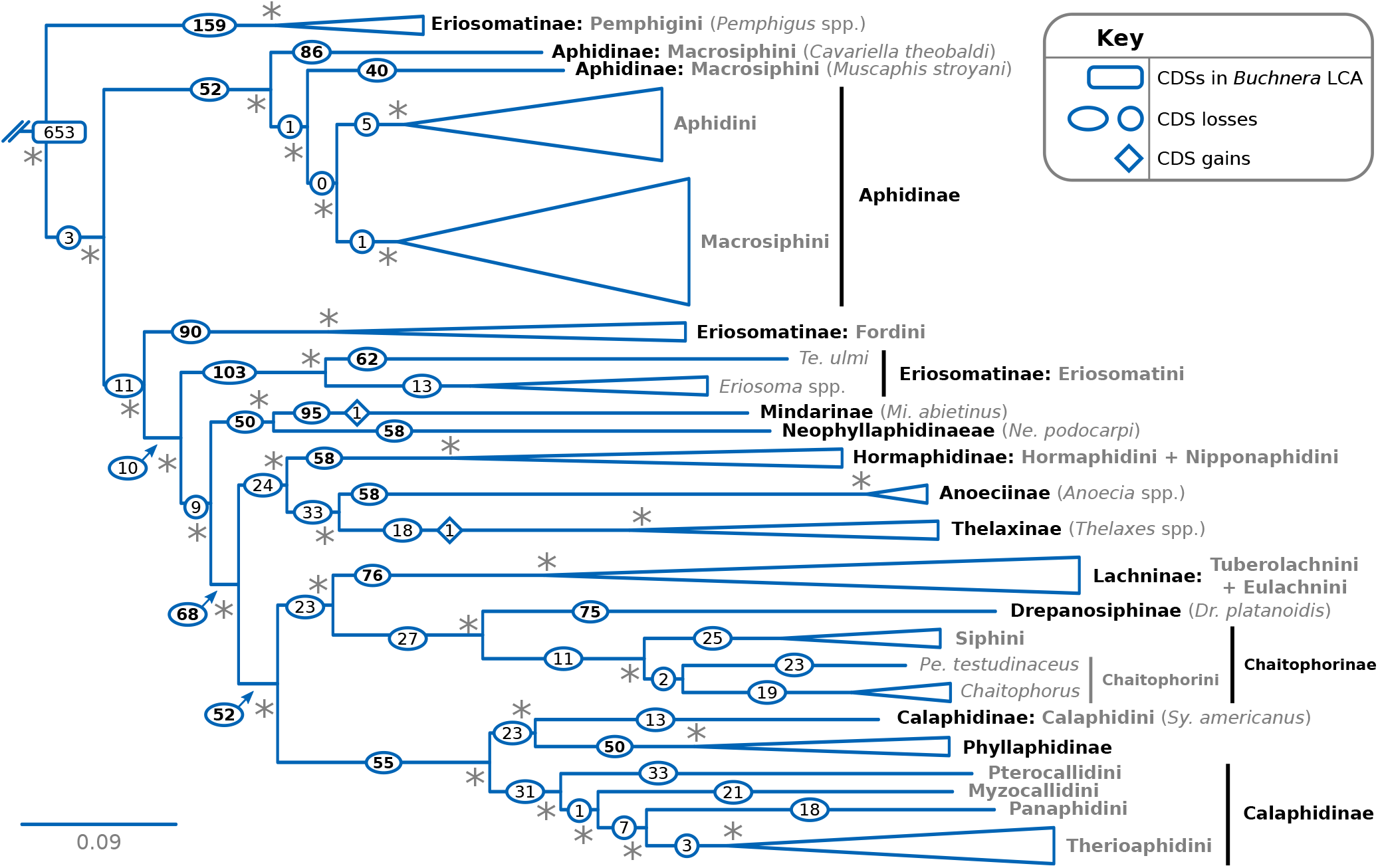
Phylogenetic relationships of *Buchnera*. Truncated Bayesian phylogenetic tree based on concatenated single-copy core proteins of selected *Buchnera*. Branches were collapsed on the basis of their aphid hosts’ subfamily classification. At the right of leaves and clades, names of aphid subfamilies and tribes are shown in bold in black and grey colours, respectively. When a subfamily is only represented by the *Buchnera* strain of one aphid species or genus, it is indicated in between parentheses.Given that *fliM*+*fliN* and *fliO*+*fliP* often occurred as fused genes, they were each counted as one protein cluster instead of two. An asterisk at nodes stands for a posterior probability of 1. *Escherichia coli*, *Pantoea ananatis*, and *Erwinia* spp. were used as outgroups. Full phylogenetic tree can be found at (supplementary figure S1, Supplementary Material online).

The phylogenetic tree, when combined with the genomic characteristics of figure 1 shows that, in agreement with Chong *et al*. (2019), the genome of *Buchnera* has undergone multiple events of genome reduction after diversification. The largest *Buchnera* genomes (*>*= 580 kbp) are retained in strains from Pemphigini, Aphidinae, Eriosomatini, Fordini, and Hormaphidinae. On the other side, small genomes of under 580 kbp have evolved once within the Eriosomatini (in *Tetraneura ulmi*), and three times in the branches leading to the Mindarinae + Neophyllaphidinae, Anoeciinae + Thelaxinae, and Lachninae + Drepanosiphinae + Chaitophorinae + Calaphidinae. After manual curation of the orthologous protein clusters of *Buchnera*, we found that the last common ancestor (LCA) of this endosymbiont coded for at least 653 proteins. Similarly to genome reduction events in *Buchnera*, marked CDS losses (*>*=40) have independently occurred in many aphid lineages. These genomic reductions are almost always accompanied by a drastic loss of protein-coding genes (CDSs). One marked exception are both *Buchnera* strains from Hormaphidinae, which hold rather large genomes (630 and 580 kbp) with a strikingly low number of CDSs (453 and 450). Surprisingly, both of these genomes keep a small number of pseudogenes (10 and 1), revealing that large portions of their genomes are devoid of any detectable gene trace (28.9% and 25.3%). When compared to the larger genomes of *Buchnera* from Aphidinae, it is becomes evident these intergenic regions originated from the pseudogenisation and eventual loss of functional genes. The LCA of both *Buchnera* from Hormaphidinae is predicted to have had at least 472 CDSs and a large genome, revealing only small CDS losses leading to each of the Hormaphidini and Nipponaphidini branches.

Most notably, we found one gene unique to each of the *Buchnera* from Mindarinae and Thelaxinae, hinting at gene acquisition in these lineages. The first gene is Tn3-family DNA-resolvase/invertase in the pYqhA plasmid of *Mi. abietinus*. This gene is similar to *Escherichia coli*’s serine recombinase PinE (prom prophage origin), which catalyzes the inversion of an 1800-bp DNA fragment (the P region), which can exist in either orientation (Plasterk and van de Putte, 1985).The second gene is a predicted amidinotransferase (FN0238 type), with best hits against Bacteroidota, present in a *Thelaxes*-specific plasmid coding for this and a *repA1* protein, similar to that present in pLeu and pYqhA plasmids (van Ham *et al*., 1997).

### Secondary symbionts complement metabolic deficiencies in six aphid lineages

Given the well-established role of *Buchnera* as an essential amino acid- and vitamin B-provider, we analysed the genes coding for enzymes involved in the biosynthesis of these nutrients across *Buchnera* (figure 3 and supplementary figure S3, Supplementary Material online). We found that *Buchnera* strains belonging to seven aphid lineages have evolved genomes potentially unable to meet the nutritional requirements of their host, displaying losses of genes namely involved in the synthesis of EAAs and B vitamins. These seven lineages include the previously reported Lachninae, *Periphyllus* spp. (Chaitophorinae: Chaitophorini), and *Si. maydis* (Chaitophorinae: Siphini), independently sequenced in this study, as well as the newly identified members of Eriosomatinae, *Anoecia* spp. (Anoeciinae), *Dr. platanoidis* (Drepanosiphinae), *Ch. stipae* (Chaitophorinae: Siphini), and *Pa. juglandis* (Calaphidinae: Panaphidini).

**Figure 3.**
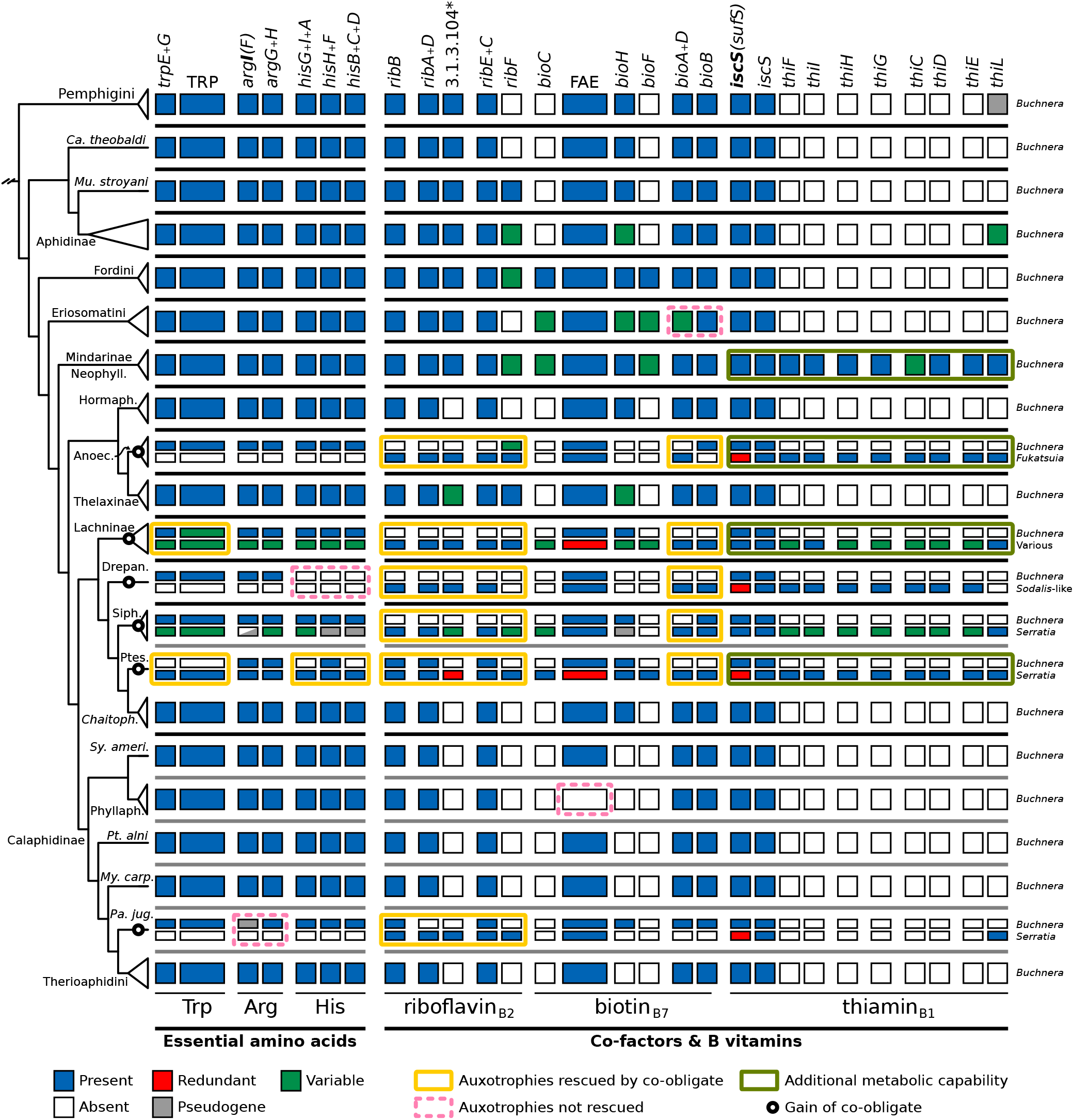
Metabolic complementarity of co-obligate symbiotic systems of aphids. Matrix summarising key metabolic capacities of obligate symbionts of different aphid species. On the left, dendrogram displaying phylogenetic relationships of *Buchnera* strains (see supplementary figure S1, Supplementary Material online). An empty circle at a node represent the acquisition of a co-obligate symbiont. At the leaves, abbreviation for aphid taxa. At the top of each box column, the name of the enzymes or pathway catalysing a reaction. At the bottom, the name of the compound synthesised by the enzymatic steps. At the right of each row of boxes, symbiont taxon name. Black and grey bars between boxes separate aphid subfamilies and tribes. TRP= anthranilate to tryptophan (*trpDCBA*), FAE= fatty acid elongation (*fab(B/F)GZ(I/V)*), Neophyll= Neophyllaphidinae, Hormaph.= Hormaphidinae, Anoec.= *Anoecia*, Drepan= *Dr. platanoidis*, Siph.= Siphini, Ptes.= *Pe. testudinaceus*, *Chaitoph.*= *Chaitophorus*, *Sy. ameri.*= *Sy. americanus*, Phyllaph.= Phyllaphidinae, *My. carp.*= *My. carpini*, *Pa. jug.*= *Pa. juglandis*. * EC number for reaction catalysed by promiscuous enzymes.

In all but one of the aforementioned aphid lineages, we were able to recover an additional symbiont genome: *Se. symbiotica* in *Periphyllus* spp., Siphini, and *Pa. juglandis*; *Fukatsuia* in *Anoecia* spp., and a *Sodalis*-like strain in *Dr. platanoidis*. Genomes of all secondary symbionts showed several symptoms of long-term association with their hosts, exhibiting clear evidence of genome reduction in comparisson to their free living counterparts. Those symbionts associated with *Anoecia* spp., *Dr. platanoidis*, and *Pa. juglandis* show the most GC-biased and reduced genomes (supplementary table S2, Supplementary Material online). On the other hand, the *Se. symbiotica* genomes of both Siphini and *Periphyllus* were more similar in size to those of facultative strains and some co-obligate ones of Lachninae (i.e. that of *Cinara tujafilina*).

Manual curation of the genes involved in the biosynthesis of EAAs and B vitamins revealed these symbiont genomes generally bear genes that are capable of rescuing auxotrophies evolved in their *Buchnera* partners. In addition, in many cases, these bacterial genomes have also lost some of the functions that are assured by *Buchnera* (figure 3), generating a clear pattern of metabolic complementarities typical of dual symbiotic systems.

In regards to B vitamins, riboflavin- and biotin-biosynthesis is often observed to be lost in *Buchnera* and being taken over/rescued by the new co-obligate symbiont. The two exceptions are in the biotin biosynthesis in *Anoecia* spp. and *Pa. juglandis*. In the former, enzymes from both symbionts are needed: *bioA*+*bioD* from the *Fukatsuia*-related symbiont and *bioB* from *Buchnera*. In the latter, the new *Se. symbiotica* symbiont has lost all three genes needed to produce this compound from KAPA, and thus *Buchnera* has retained this role. As reported for other *Periphyllus* spp. by Monnin *et al*. (2020), we corroborate the tryptophan- and histidine-biosynthetic role of the *Se. symbiotica* endosymbiont in the newly sequenced genome from *Pe. testudinaceus*. Nonetheless, we did not observed the phenylalanine biosynthetic role being lost in *Buchnera* from *Pe. testudinaceus*. Upon closer inspection of the *Buchnera* draft genomes and annotations of *Periphyllus* spp. analysed by Monnin *et al*. (2020), we found the presence of a conserved poly(A) region across *Buchnera* from *Periphyllus* spp. in the *pheA* gene. Due to the conserved nature of this region, we considered it as likely rescued by transcriptional frameshifting, as this rescue mechanism has been experimentally demonstrated in other *Buchnera* and *Blochmannia* strains (Tamas *et al*., 2008) and shown to be maintained by natural selection in the latter (Wernegreen *et al*., 2010). In addition, similar poly(A) regions disrupting the *pheA* gene are present in all other sequenced *Buchnera* strains from Chaitophorinae. We found the thiamin-biosynthetic capacity (vitamin B_1_) to be largely retained in the symbiotic consortia from Mindarinae, Neophyllaphidinae, Anoeciinae, Lachninae, Drepanosiphinae, Siphini, and *Pe. testudinaceus*. Most surprisingly, in the case of Mindarinae we found the retention of thiamin biosynthetic genes in newly sequenced *Buchnera* from Mindarinae (all) and Neophyllaphidinae (no *thiC*). The genes are present in four syntenic regions: between the *purH* and *rpoC* (*thiCEFSGH*), *dcd* and *metG* (*thiD*), *ribE* and *ribD* (*thiL*), and *dxs* and *yajR* (*thiI*). The location of the *thi* genes is syntenic with *Erwinia* spp., suggesting these genes were present in the LCA of *Buchnera* and have been repeatedly lost. Supporting this hypothesis, we observed the presence of a *thiL* gene/pseudogene in *Buchnera* from Pemphiginae and many Aphidinae, with top BLASTp hits *vs.* NCBI’s nr to *Erwnia* and *Pantoea* bacteria, which are the closest free-living relatives of *Buchnera* (Husník *et al*., 2011). Finally, *Buchnera* from Phyllaphidinae have lost the genes involved in fatty acid elongation, which would make *Buchnera* dependant on the host for the production of its own membrane. However, as is the case for other strains, it would be able to synthesise biotin from KAPA.

Regarding essential amino acids, we corroborated the collaboration of both symbiotic partners in the production of tryptophan in some Lachninae (Lamelas *et al*., 2011b; Manzano-Marín *et al*., 2016) as well as the takeover of this role and the histidine biosynthesis by the co-obligate *Se. symbiotica* in *Periphyllus* spp. (Monnin *et al*., 2020). In addition, we uncovered the complete loss of histidine-biosynthetic genes in *Buchnera* from Drepanosiphinae and a loss of the *argI* gene, rendering *Buchnera* and potentially the aphid host dependant on histidine and/or L-citruline for the production of these two essential amino acids.

### Aphids are associated with a large and diverse pool of secondary symbionts

Most known co-obligate symbiotic lineages have facultative counterparts (Manzano-Marín *et al*., 2018, 2020; Meseguer *et al*., 2017), and thus, exploring the symbiotic microbiota from a large diverse aphid pool can provide clues into the available potential source for new co-obligate symbionts. Our 16S rRNA amplicon survey of 223 aphid samples yielded ~5.4 million sequencing reads with an average of ~11,300 reads per aphid sample kept. Those reads were distributed into 287 clusters, with most of these assigned to *Buchnera aphidicola* and nine known facultative or obligate symbionts of aphids (figure 4 and supplementary figure S2, Supplementary Material online), including the recently described *Erwinia haradaeae* (Manzano-Marín *et al*., 2020), *Fukatsuia* in some *Cinara* spp. (Manzano-Marín *et al*., 2017; Meseguer *et al*., 2017), and *Bacteroidota* which has been found replacing *Buchnera* in *Geopemphigus* spp. (Chong and Moran, 2018). In addition, some clusters were designated as *Acetobacteraceae*- and *Gillamella*-related bacteria, known as gut-associated symbionts of diverse insects (Brown and Wernegreen, 2019; Holley *et al*., 2022; Kwong and Moran, 2013; Pais *et al*., 2018; Smith and Newton, 2020). The remaining reads were assigned to ubiquitous bacteria that could be of an environmental source and not representative of the aphid microbiota. In fact, these bacterial taxa were not always found across PCR replicates of the same sample and were sometimes found in the controls (supplementary figure S2, Supplementary Material online). From the bacterial taxa, *Se. symbiotica* was the most common secondary endosymbiont detected in our sampling, being present in 77 out of 223 samples and across seven subfamilies. In addition to *Se. symbiotica*; *Hamiltonella*, *Wolbachia*, *Fukatsuia*, and *Sodalis*-like bacteria were found in 23 (across five subfamilies), 22 (across five subfamilies), 18 (across seven subfamilies), and 15 samples (across six subfamilies), respectively.

**Figure 4.**
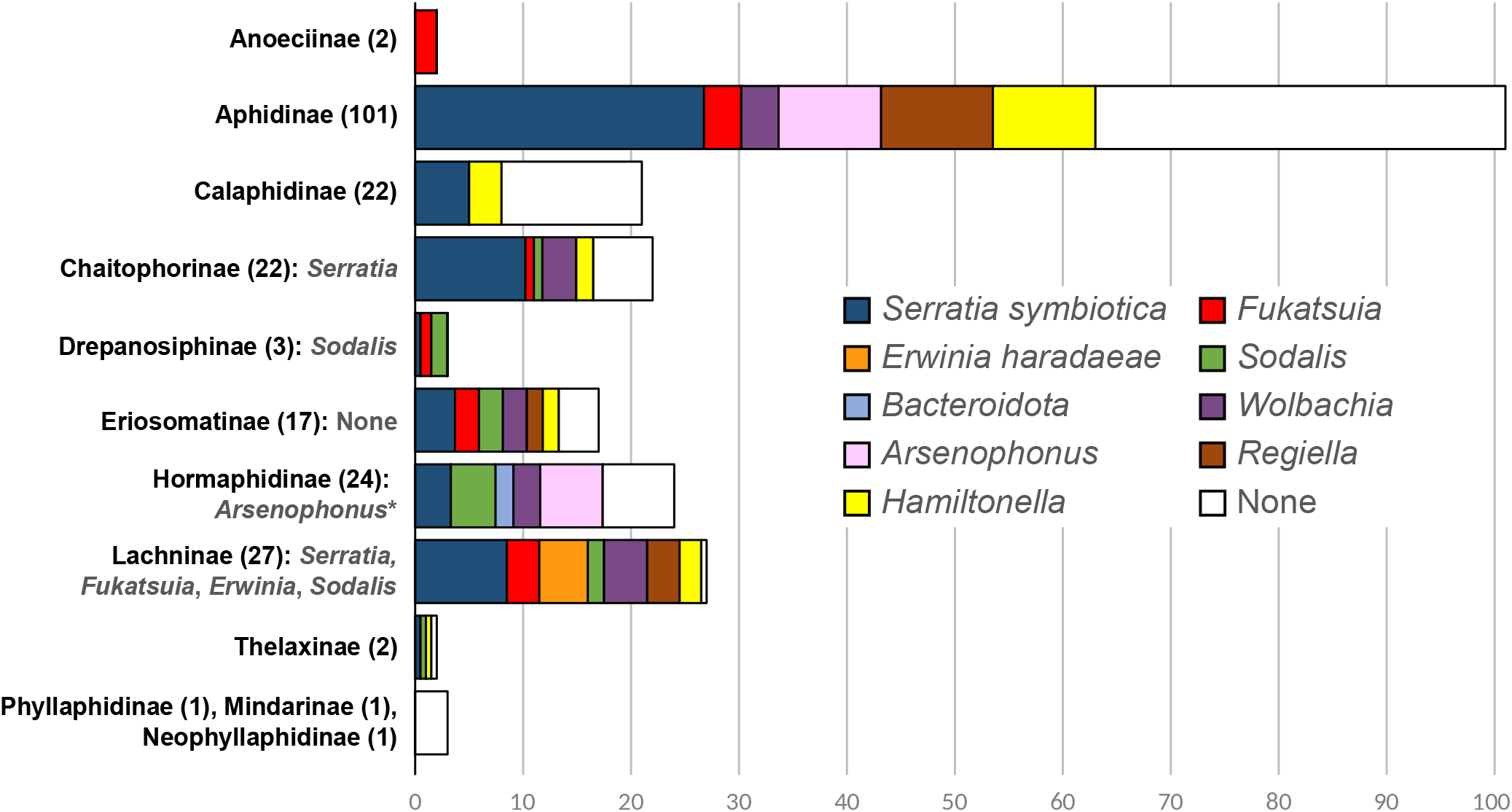
Secondary symbiont diversity across aphid subfamilies. Stacked barplot displaying the scaled absolute count of symbiont taxa found across samples of specific subfamilies. To the left of the bars, the subfamily name is indicated followed by the number of samples included in this survey between parentheses. Family names and counts are followed by the putative and known co-obligate symbiont taxa as inferred from the present study or previous genomic analysis studies. * Recently published by Yorimoto *et al*. (2022)

This 16S amplicon survey is in agreement with our analysis of genomic data: when a co-obligate association has been inferred in an aphid species by genome-based metabolic inference, the symbiont taxon was found in all samples of that species and often in closely related ones. For example, we detected the presence of *Se. symbiotica* in all species of *Periphyllus* spp. (Chaitophorinae: Chaitophorini), where it has previously been reported as a co-obligate symbiont (Monnin *et al*., 2020). This symbiont was also detected in Siphini from our sampling (Chaitophorinae: Siphini), as well as in various Lachninae species, where it is known as an obligate partner of many species (Lamelas *et al*., 2011b; Manzano-Marín and Latorre, 2014; Manzano-Marín *et al*., 2016, 2017; Meseguer *et al*., 2017). The description of the dual symbiotic system of *Si. maydis* has been very recently independently reported by Renoz *et al*. (2022b), which resulted from parallel sequencing efforts and showing similar findings to ours for this system. *Erwinia* was also found in all samples from a specific clade of *Cinara* (Lachninae: Eulachnini), validating its co-obligate status (Manzano-Marín *et al*., 2020). In addition, we found *Se. symbiotica*, *Fukatsuia* and a *Sodalis*-like bacterium in all samples of *Pa. juglanidis*, Anoeciinae and Drepanosiphinae aphids, respectively.

Our analysis also corroborated the absence of *Buchnera* in the one sample of *Cerataphis brasiliensis* (Hormaphidinae: Cerataphidini), a species in which the ancient obligate symbiont has likely been replaced by a ”Yeast-like symbiont” (Vogel and Moran, 2013). Further, although a *Bacteroidota* symbiont has been found replacing *Buchnera* in a member of the Fordini (*Geopemphigus* spp.; Chong and Moran, 2018), we did not recover any secondary bacterial 16S rRNA sequence belonging to this bacterial taxon in the Fordini sampled in this study (*Baizongia pistaciae*, *Forda* sp., and *Geoica* sp.). In fact, 16S rRNA sequences assigned to *Bacteroidota* were only found in two samples of Hormaphidinae aphids in our survey (*Astegopteryx bambusae* ACOE3753 and *Reticulaphis mirabilis* ACOE3769). *Arsenophonus* was found in many samples of Hormaphidinae, where it has been previously found to be common within this subfamily and, very recently, even found to be establishing a co-obligate association in *Ceratovacuna japonica* (Hormaphidinae: Cerataphidini; Yorimoto *et al*. 2022). Aphidinae hosted many secondary symbionts, however, none appear to be fixed at least at the tribe or even genus level.

### Newly identified co-obligate symbionts have repeatedly evolved from within well-known symbiotic lineages

Phylogenetic placement of the newly sequenced *Serratia*-related co-obligate symbionts confirmed their taxonomic assignment as *Se. symbiotica* (figure 5A). The recovered phylogenetic tree suggests at least two independent origins of the co-obligate *Se. symbiotica* endosymbionts in *Periphyllus* spp., with support for a single origin of this symbiont lineage in the LCA of *Pe. aceris* and *Pe. acericola*. Additionally, the resulting phylogeny does not support a common origin for the co-obligate symbionts of Siphini aphids, suggesting repeated replacements within this tribe. Regarding the co-obligate symbionts of *Anoecia* spp., they are recovered as a sister group to *Fu. symbiotica*. The 16S identity of the *Fukatsuia*-like co-obligate symbionts of Anoeciinae aphids falls below the recommended species threshold of 98.7% (95.79% and 96.04%; Chun *et al*., 2018), but above the recommended minimum threshold for being classified as the same genus (Yarza *et al*., 2014). Nonetheless, both *Anoecia* spp. symbionts show a 16S sequence identity of 98.77%, suggesting they belong to the same molecular species. Co-obligate *Sodalis*-like symbionts have been identified in several aphid species (Manzano-Marín *et al*., 2017, 2018). The newly identified *Sodalis*-like co-obligate symbiont of *Dr. platanoidis* does not cluster with any of the previously identified aphid *Sodalis*-like symbionts (figure 5B). In fact, the tree suggests most co-obligate *Sodalis*-like endosymbionts from aphids have evolved independently, with those of *Eulachnus* and *Cinara (Schizolachnus)* spp. having likely evolved from closely related ancestor strains.

**Figure 5.**
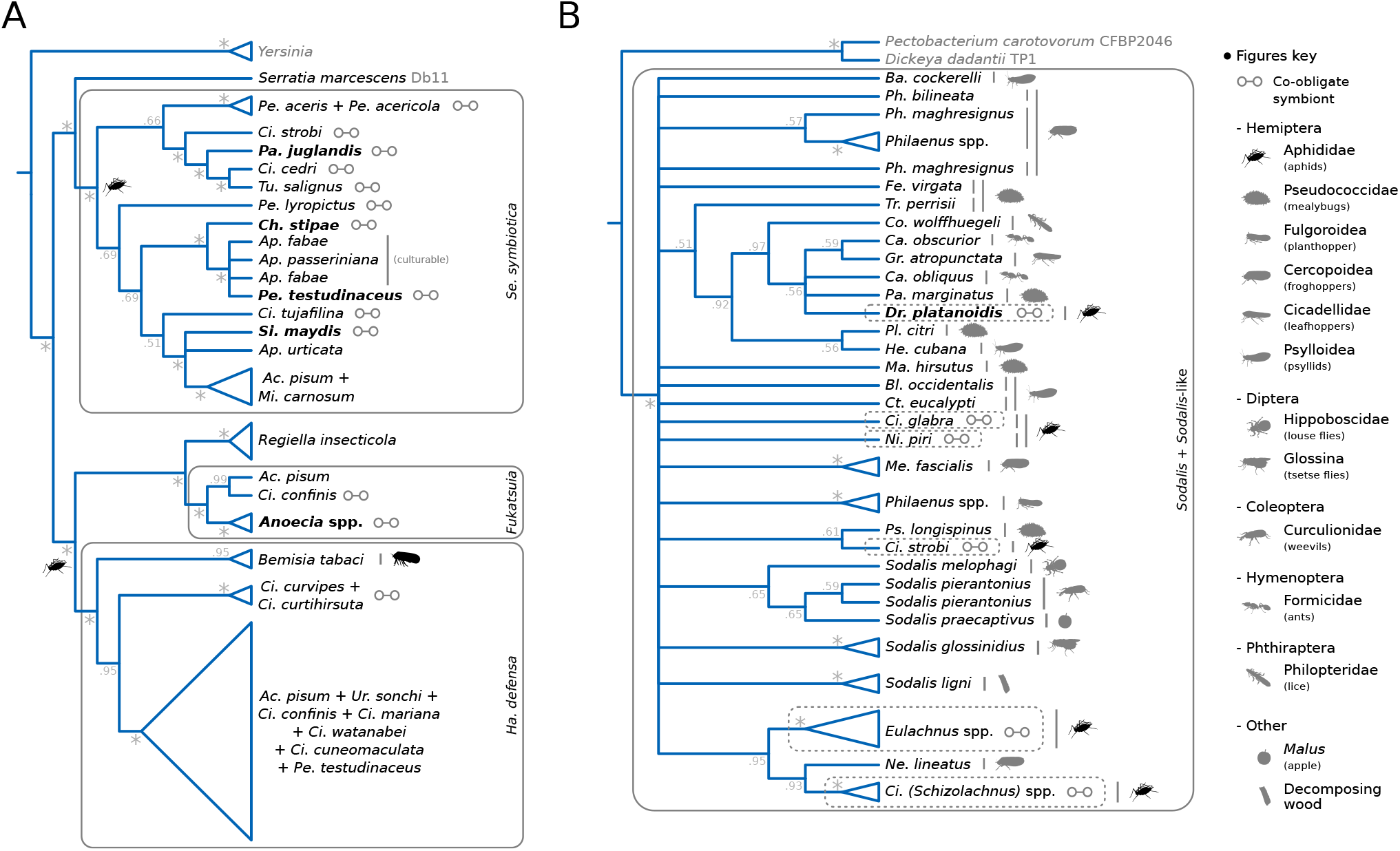
Phylogenetic relationships of aphid co-obligate endosymbionts. **(A)** Phylogenetic placement of *Yersinia*- and *Serratia*-related co-obligate endosymbionts. Silhouettes at nodes or at leaves indicate host relationship of clade. Grey boxes with binomial names on the side indicate aphid symbiotic taxa of interest. **(B)** Phylogenetic placement of *Sodalis*-related co-obligate endosymbionts. Grey box delimits the *Sodalis* + *Sodalis*-related clade. Silhouettes at branch ends indicate identity of host relationship of leaf/clade. Dotted boxes highlight *Sodalis*-like symbionts of aphids. For both A and B, abbreviated nomenclature is used when referring to the symbiont’s host name. Names in bold, highlight newly sequenced endosymbiont lineages. Names in grey mark taxa used as outgroups.

### Co-obligate symbionts in *An. corni* and *Si. maydis* reside inside bacteriocyte cells

In order to investigate the distribution of the newly identified co-obligate endosymbionts inside of their aphid hosts, we analysed available preserved specimens of *An. corni* and *Si. maydis* embryos through FISH miscroscopy using specific probes for each symbiont (figure 6). We corroborated and taxonomically identified the secondary symbionts previously observed by transmission electron microscopy by Michalik (2010); Michalik *et al*. (2014). Taken together, these observations support that the *Fukatsuia* symbiont of *An. corni* resides in separate bacteriocytes to those of *Buchnera* and shows a spherical shape. Similarly, we observed the *Se. symbiotica* symbiont of *Si. maydis* inhabiting different bacteriocytes to those of *Buchnera*. However, the co-obligate endosymbiont shows an elongated rod-shape, more typical of facultative and early co-obligate symbionts (Manzano-Marín and Latorre, 2014; Moran *et al*., 2005).

**Figure 6.**
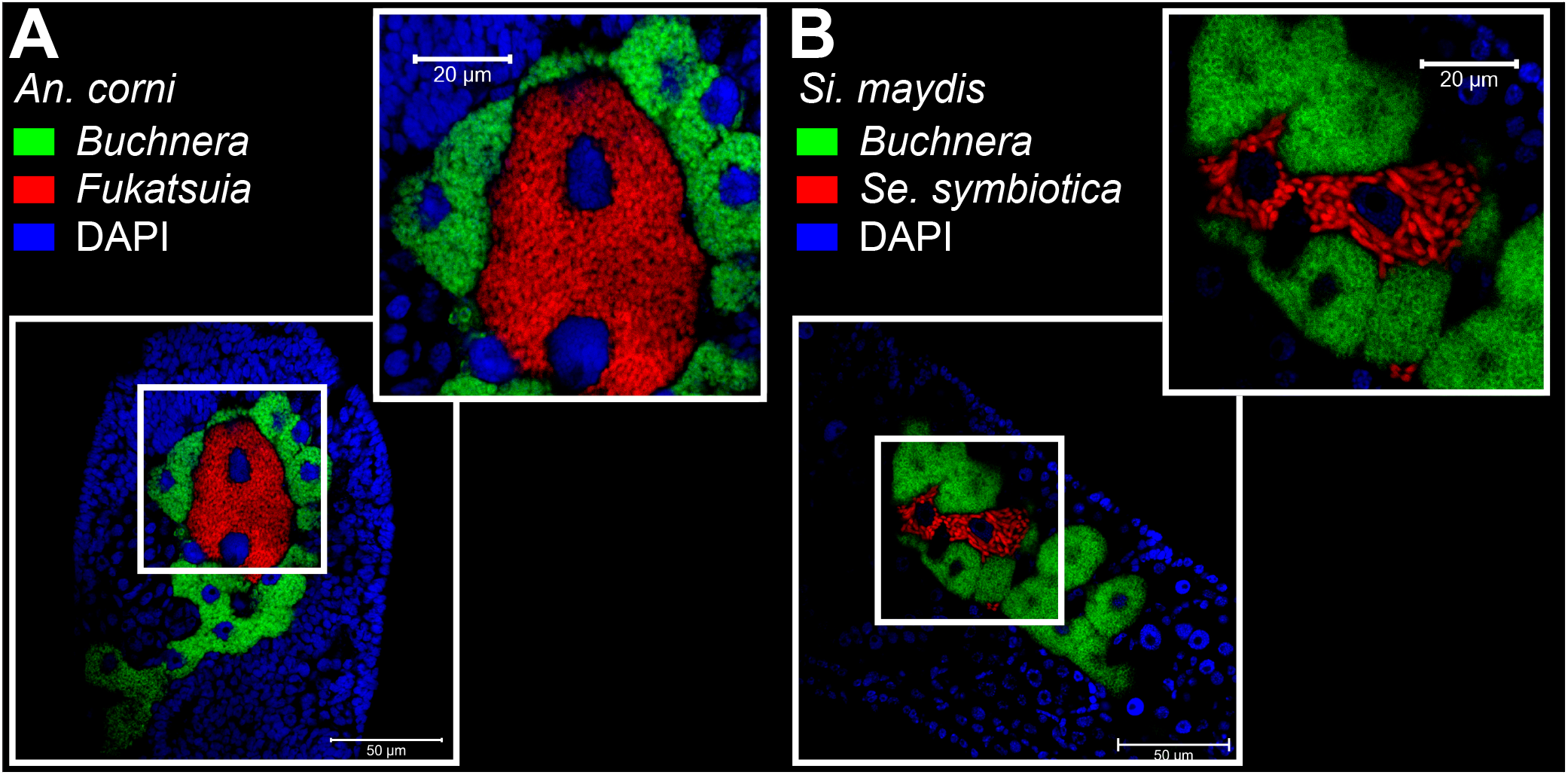
Location and morphology of co-obligate symbionts in selected aphid species. Merged FISH microscopic images of aphid embryos from **(A)** *An. corni* (lateral) **(B)** *Si. maydis* (tilted lateral). Co-obligate symbionts’ signal is shown in red, *Buchnera*’s in green, and DAPI’s (staining DNA, highlighting host nuclei) in blue. Thick white boxes indicate the magnified region depicted in the top-right of each panel. The scientific name for each species along with the false colour code for each fluorescent probe and its target taxon are shown at the top-left of each panel. Unmerged images can be found in https://doi.org/10.5281/zenodo.6394197.

### ’*Candidatus* Fukatsuia anoeciicola’ sp. nov

’*Candidatus* Fukatsuia anoeciicola’ (an.oe.ci.i’co.la. N.L. fem. n. *Anoecia*, an aphid genus; L. masc./fem. n. suff. - cola, inhabitant, dweller; N.L. fem. n. *anoeciicola*, inhabiting *Anoecia* aphids).

We propose the specific name ‘*Candidatus* Fukatsuia anoeciicola’ for the monophyletic lineage of enterobacterial endosymbionts from the ‘*Candidatus* Fukatsuia’ genus hitherto exclusively found affiliated as co-obligate nutritional symbionts in *Anoecia* species (Hemiptera: Aphididae: Anoeciinae). In embryos of *Anoecia corni*, ‘*Candidatus* Fukatsuia anoeciicola’ is found co-inhabiting the bacteriome intracellularly in different bacteriocytes to those of *Buchnera aphidicola* (figure 6; Michalik *et al*., 2014). In oviparous *An. corni*, ‘*Candidatus* Fukatsuia anoeciicola’ has a spherical shape (mean cell diameter of 2.08µm), it is located intracellularly in bacteriocytes surrounded by those of *Buchnera*, and can also be found in oocytes (Michalik *et al*., 2014).

## Discussion

Long-term associations with strict vertically-transmitted bacteria often results in drastic genome reduction of the bacterial symbiont, which can sometimes lead to impairment of the symbiotic functions (*e.g.* host’s nutrition) (McCutcheon *et al*., 2019). In this context, replacement or complementation of the primary symbiont with a new one can maintain the stability of the symbiotic system. There are now many descriptions of symbiont replacement and complementation (Sudakaran *et al*., 2017), in sap feeding insects, blood-feeding ticks and lice (Buysse *et al*., 2021; Říhová et al., 2021), and weevils (Conord *et al*., 2008; Lefèvre *et al*., 2004; Toju *et al*., 2013). The nutritional association of *Buchnera aphidicola* with aphids has persisted for millions of years and has widely been regarded, until very recently, as mostly existing in one-to-one obligate associations with its host, in comparison to nutritional symbioses observed in other hemipterans. While both early and recent microscopic investigations into diverse aphids has revealed several cases of likely dual symbiotic systems (Buchner, 1953; Fukatsu, 2001; Fukatsu and Ishikawa, 1993; Michalik, 2010; Michalik *et al*., 2014), little is known about the diversity and nature of these associations. Here we add to the recent evidence that the aphid-*Buchnera* evolutionary history is marked by repeated losses of essential metabolic capacities and complementation by a new co-obligate partner. We show that 12 aphid species (8 sequenced by us) from six subfamilies in which *Buchnera* has lost one or several essential symbiotic functions. This study, confirms recent observations in Chaitophorinae on two different genera, *Periphyllus* and *Sipha* (Monnin *et al*., 2020; Renoz *et al*., 2022b). Our study shows that *Se. symbiotica* also occurs as a co-obligate nutritional symbiont in the related *Chatosiphella* genus. However, we show that *Buchnera* from the closely related *Chaitophorus* genus preserves an intact repertoire to synthesise all EAAs and B vitamins. Along the same line, very recently published evidence strongly suggests that members of the *Ceratovacuna* genus (Hormaphidinae: cerataphidini) rely on two nutritional symbionts, *Buchnera* and *Arsenophonus* (Yorimoto *et al*., 2022). According to our analyses, representative from the other two tribes of Hormaphidinae, do not depend on additional symbionts from nutritional complementation. Thus, in both Hormaphidinae and Chaitophorinae, losses of symbiotic functions happened after the diversification of the subfamily, and in the latter, after diversification of the tribe Chaitophorini. In most cases, *Buchnera*’s auxotrophies are rescued by a new bacterial partner, with the marked exceptions of two Eriosomatini (biotin), Drepanosiphinae (histidine), and Calaphidinae (Arginine), for which horizontal gene-transfer event to the host might play a role (see below for expanded discussion).

We found *Se. symbiotica* associated as the new co-obligate partner in now four subfamilies with at least four independent acquisitions (figure 7). However, as for Lachninae aphids (Manzano-Marín *et al*., 2017; Meseguer *et al*., 2017), repeated replacement of co-obligate endosymbionts has likely occurred in *Periphyllus* and Siphini aphids: the *Se. symbiotica* strains from these aphids do not form a well-supported monophyletic clade (figure 5). In addition, repeated replacement has likely also occurred within Drepanosiphinae: while we found *Drepanosiphum* and *Drepanosiphoniella* hosted *Sodalis*-like symbionts, *Yamatocallis* spp. have been shown to harbour unrelated gammaproteobacterial obligate symbionts (Fukatsu, 2001). These patterns of repeated symbiont replacement would thus be similar to those observed in mealybugs (Husnik and McCutcheon, 2016), psyllids (Sloan and Moran, 2012), and Auchenorrhyncha (Bennett and Moran, 2013), where the primary obligate endosymbiont is retained, but the secondary co-obligate one shows less stability over evolutionary time.

**Figure 7.**
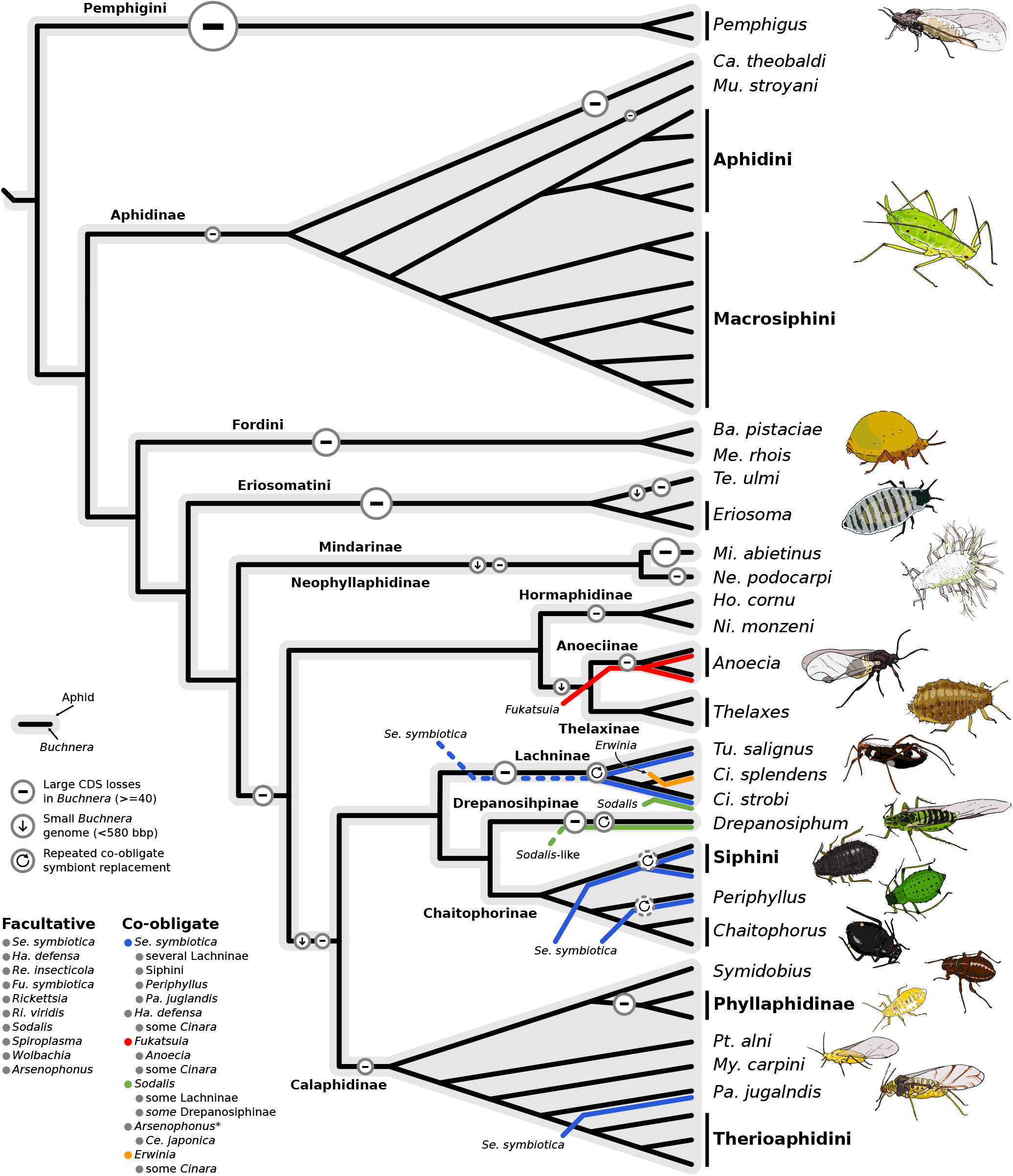
Evolutionary history of co-obligate symbioses in Aphididae. Diagram summarising results obtained from the genomic analysis presented in this work. Cladogram was made using the reconstructed phylogeny for *Buchnera* symbionts. Black lines in the cladogram represent *Buchnera*. Coloured lines are used to represent the acquisition of co-obligate symbionts. Dotted lines represent uncertainty in the acquisition of a specific taxa. Dotted circles represent uncertainty in symbiont replacement dynamics within the aphid lineage. On the right, aphid cartoons depicting variety of species across the Aphididae.

*Se. symbiotica* was the most common bacterial lineage detected in our 16s rRNA survey, confirming its predominance as a natural facultative symbiont of aphids (Pons *et al*., 2022; Zytynska and Weisser, 2016). In fact, *Se. symbiotica* has not only been recorded in aphids, but has also been identified as a possible facultative endosymbiont in Japanese populations of the hemlock woolly adelgid *Adelges tsugae* (von Dohlen *et al*., 2013). Strains of this symbiont have been recorded as protecting against heat-stress (Chen *et al*., 2000; Montllor *et al*., 2002; Russell and Moran, 2006) and providing resistance against parasitoid wasps (Oliver *et al*., 2003). In fact, strain IS has and even been shown to compensate, under laboratory conditions, for the elimination of the primary obligate endosymbiont *Buchnera* in the pea aphid *Acyrthosiphon pisum*, albeit for only 24 generations (Koga *et al*., 2003) (suggesting some functional redundancy with *Buchnera*). Given the multiple benefits conferred by this endosymbiont, the potential for horizontal transmission through the host plant of the aphid host (Pons *et al*., 2019), and the capacity that some strains might have to take-over (at least) part of the symbiotic contributions of *Buchnera*, it is not unexpected to find it widespread across aphid taxa and populations, where it might easily become more common when the right environmental and biological conditions arise. The pervasiveness of *Se. symbiotica* strains in aphid populations has probably facilitated their adaptation as new co-obligate partners upon *Buchnera*’s gene losses. However, *Se. symbiotica* is not the only bacterial lineage that has become essential to its aphid hosts. A *Fukatsuia*-related symbiont (*Fukatsuia anoeciicola*) was revealed as a co-obligate partner of Anoeciine aphids. Another species of this symbiotic lineage, *Fukatsuia symbiotica*, has already been reported as a co-obligate endosymbiont of Lachninae (Manzano-Marín *et al*., 2017; Meseguer *et al*., 2017). Our 16S rRNA amplicon survey revealed that *Fukatsuia*-related symbionts are found in populations of aphid species from different subfamilies, but much less frequently than *Se. symbiotica*. *Fukatsuia* symbionts have been associated with several benefits to the pea aphid (i.e resistance against parasitoid wasps, fungal pathogens, and increased reproduction after heat-stress), however only when co-infecting with other symbiont species (Donald *et al*., 2016; Doremus and Oliver, 2017; Heyworth and Ferrari, 2015). Additionally a *Sodalis*-like symbiont was identified as a co-obligate partner of Drepanosiphinae. Despite *Sodalis*-related strains being widely found across arthropods, they have been rarely reported in aphids. Some of the few strains found in aphids have been described as being in an obligate association with some Lachninae, mainly *Cinara* spp. (Manzano-Marín *et al*., 2017, 2018; Meseguer *et al*., 2017). We have nonetheless identified this symbiont occasionally in six subfamilies. With limited knowledge on *Sodalis* and *Fukatsuia* potential benefits/costs as facultative symbionts, it is hard to speculate on the environmental conditions favouring their expansion in populations, and thus, their adoption as a co-obligate partners. In any case, our data on *Sodalis*, along with the uncovering of *Erwinia haradaeae* as secondary co-obligate symbiont in some *Cinara* spp. and the *Bacteroidota* symbiont replacing *Buchnera* in *Geompemphigus* spp., shows that, even relatively rare symbionts (in extant populations) and environmental bacteria not known as symbionts of the host, can eventually become essential to it. These patterns mirror what has been observed in other sap- and blood-feeding lineages hosting dual symbiotic systems. For example, in whiteflies, a widespread but rare arthropod symbiont, *Arsenophonus*, has evolved to complement its degenerated *Portiera* symbiont. Conversely, phylogenetic placement of tick symbionts suggests they have evolved from common pathogens found in blood meals (Duron *et al*., 2018). Hence, our study adds to the current evidence that obligate symbionts can derive not only from facultative beneficial symbionts but also sometimes from less common bacterial symbionts as well as strains that occur in the environment (Hosokawa *et al*., 2016).

Among the very speciose Aphidinae subfamily, which shelters a large diversity of facultative symbionts (figure 4), and where 56% of the samples were positive for at least one facultative symbiotic taxon, we found no evidence of putative co-obligate endosymbionts, including in the banana aphid *Pentalonia nigronervosa*, corroborating previous genomic re-analyses (Manzano-Marín, 2020). As previously mentioned, nutritonal co-obligate associations have been suggested for two Aphidinae species (Monnin *et al*., 2020). However, extensive review of this work casts important doubts regarding this conclusion (Manzano-Marín, 2020).

A common pattern in the dual nutritional symbioses revealed here is the riboflavin biosynthetic-role takeover by the new co-obligate symbiont. Biotin was the second most common nutrient predicted to be supplied by the secondary co-obligate endosymbionts: it was found in all cases except in the co-obligate association observed in *Pa. juglandis*. A similar vitamin-biosynthetic role for the new co-obligate endosymbionts can be found in some psyllids (Sloan and Moran, 2012) as well as some adelgids (Dial *et al*., 2022; Szabó *et al*., 2022). In contrast, in other adelgids, retention of the B vitamin biosynthetic genes can also be observed in the primary symbionts, and not in the newly evolved symbiotic partners. In mealybugs, genes from these pathways can even be found missing in all symbiont genomes, while being found either partially (riboflavin) or entirely (biotin) in the hosts’ genomes, originating from horizontal gene-transfer events (Husnik and McCutcheon, 2016). Therefore, this pattern seems to be system specific, and likely dependent on the background of the previously existing primary obligate endosymbiont. To our knowledge, no information is available on the specific life stage on which the aphid might depend on this nutrient; which precludes elaborating scenarios on how this pathway can be lost in *Buchnera*. In addition, thiamin was found to be retained in different lineages of co-obligate symbionts with a reduced genome, while lost in others as well as in all *Buchnera* genomes but the ones associated with Mindarinae and Neophyllaphidinae (figure 3). Consequently, complete and almost-complete pathways (lacking the thiamine-monophosphate kinase gene *thiL*) are found in obligate symbiotic systems of five subfamilies. The retention of this pathway in highly reduced genomes of endosymbionts suggest an important role for this nutritional role. This has been previously proposed in the endosymbiotic systems of a monophyletic group of *Cinara* spp., where the thiamin-biosynthetic genes have been serially horizontally transferred from a *Sodalis*-like bacteria to the now co-obligate *Er. haradaeae*, and further to a tertiary co-obligate *Ha. defensa* (Manzano-Marín *et al*., 2020). Interestingly, the *Buchnera* strains that retain this pathway are associated exclusively with aphids feeding on conifers, as *Cinara* spp. Sap feeders are generally assumed to have very similar nutritional requirements, relying exclusively on a sugar rich diet that is deficient in EAAs and vitamins. However, based on the plant taxon they feed on, phloem sap might vary in their vitamin and amino acid abundance. Unfortunately, and to our knowledge, no comparative phloem sap analysis of conifers *vs.* angiosperms is available which could aid in inferring whether this symbiotic function can be related to host plant association. Further, other di-symbiotic systems maintaining thiamin genes analysed here are associated with more diverse botanical families (i.e. some herbaceus plants for *Si. maydis*). However, the presence of this pathway in the *Se. symbiotica* of *Si. maydis* (Siphini) can be related to a recent acquisition of this co-obligate strain: its genome size remains similar to that of facultative strains associated, for instance, with *Ac. pisum* (Burke and Moran, 2011; Nikoh *et al*., 2019). Therefore, the retention of this pathway might not necessarily be related to any essential nutrient supplementation role. More studies elucidating the importance of thiamin during aphid development and its abundance in the phloem-sap of diverse host plant families are necessary to fully understand the evolutionary processes leading to its losses and gains throughout the diversification of aphids.

The amino acid-biosynthetic genes are almost always retained in *Buchnera*, while lost in the secondary co-obligate symbionts (figure 3). Marked exceptions are those of the split pathway between the co-obligate symbionts of the Lachninae aphids *Cinara cedri* and *Tu. salignus* (Lamelas *et al*., 2008; Manzano-Marín *et al*., 2016) as well as the takeover of the biosynthesis of this compound plus histidine in *Periphyllus* spp. (Monnin *et al*., 2020; Renoz *et al*., 2022b). Most surprising was the identification of evolved auxotrophies in *Buchnera* that were not rescued by the co-obligate symbionts retrieved in our dataset. This was the case for arginine in *Pa. juglandis*, histidine in *Dr. platanoidis*, fatty acid elongation in Phyllaphidinae, and biotin in *Er. grossulariae* and *Te. ulmi*. We hypothesise that either the aphid is able to supplement precursors through horizontally transferred genes or the specific diet of the aphid is able to rescue these auxotrophies. In fact, horizontally transferred genes from diverse bacteria have been predicted and/or shown to support nutrient biosynthesis in Pseudococcinae (Husnik and McCutcheon, 2016; Husnik *et al*., 2013; Szabó *et al*., 2017), psyllids (Sloan *et al*., 2014), and the whitefly *Bemisia tabaci* (Bao *et al*., 2021; Luan *et al*., 2015; Ren *et al*., 2020; Xie *et al*., 2018). In particular, certain genes whose product are involve in the biosynthesis of arginine and biotin have been shown to be acquired by diverse hemipterans through horizontal gene-transfer (HGT) and hypothesised to be involved in complementing their nutritional symbiont’s biosynthetic pathways (Husnik and McCutcheon, 2016; Luan *et al*., 2015). Another scenario would be the presence of an additional symbiont not detected by our genomic sequencing. In this vein, it is important to note that the presence of a *Fukatsuia* strain was detected in both *Dr. platanoidis* samples, by 16S rRNA amplicon sequencing (supplementary table S2, Supplementary Material online). However, in the case of both *Dr. platanoidis* and *Pa. juglandis*, microscopic studies have revealed the presence of only two different bacteria (Buchner, 1953). Additionally, a *Sodalis*-like strain was also detected in the related *Drepanosiphoniella fugans*, and while no deeper investigation of this genus/species was performed, it hints to a possible ancestral association with a *Sodalis*-like lineage. It is noteworthy to mention that both currently available sequenced strains of *Fukatsuia* symbionts do not preserve any genes in the histidine biosynthetic pathway (Patel *et al*., 2019). Similarly, we observed a lack of these genes in the assembled genome for the *Fukatsuia* strain associated with *Dr. platanoidis*. Lastly, both *Sodalis*-like and *Se. symbiotica* symbionts of the aforementioned species show typical characteristics of well established obligate vertically transmitted symbionts: a lower G+C content and smaller genome size than their facultative/free-living symbiotic counterparts as well as clear metabolic complementarities with their corresponding symbiotic partner. Therefore, currently available evidence points towards these endosymbiotic systems being made up of *Buchnera* and one additional secondary co-obligate symbiont.

Our study also revealed patterns of *Buchnera* degradation: this primary symbiont has undergone multiple events of CDS losses, which accordingly is most commonly accompanied by a reduction in genome size. While the acquisition of co-obligate endosymbionts is often associated to a drastic gene loss (figure 7), there are also some exceptions to this pattern (figure S4, Supplementary Material online). In Calaphidinae and Phyllaphidinae, *Buchnera* are generally below 450 Mb (as small as in Lachninae) but this reduction was not concomitant to dependency on a new partner, as gene losses did not affect symbiotic functions. On the other hand, very few genes are actually lost in the branch leading to *Pa. juglandis* (18, out of which 8 are related to essential amino acid- and B vitamin-biosynthesis). A similar case is observed in the breanch leading to *Periphyllus* spp. This shows that it is the inactivation of very few functional genes that actually triggers dependency on an additional symbiont and not necessarily large-scale genome decay. We also observed large genomes with a low coding density in Hormaphidinae aphids. Such large amounts of ”deserted” DNA are more commonly observed in some transitional symbiont genomes (Manzano-Marín and Latorre, 2016; Manzano-Marín *et al*., 2018). To our knowledge, such low coding densities in such a small symbiont genome has only been reported for the primary obligate endosymbiont of mealybugs, *Tremblaya princeps* (McCutcheon and von Dohlen, 2011), as well as for several of their co-obligate *Sodalis*-related endosymbionts (Husnik and McCutcheon, 2016). This suggests that, although rarely observed, large gene losses can precede genome reduction, even in a small compact symbiont genome. More surprisingly, we found the recent acquisition of genes to *Buchnera* plasmids in *Mi. abietinus* and *Thelaxes* spp. The role these two newly gained genes might play are quite different. The DNA-resolvase/invertase found in the pYqhA plasmid of the *Buchnera* from *Mi. abietinus* might play a role in promoting inversions within this plasmid. On the other hand, the predicted amidinotransferase carried by *Buchnera* from *Thelaxes* spp. might play a yet unknown metabolic role. Regarding the DNA-resolvase/invertase, it keeps similarity to the PinE protein of *E. coli* which is responsible for a naturally occurring inversion of the so-called P-region (Plasterk and van de Putte, 1985). In fact, historical inversions are observable among pLeu plasmids of *Buchnera* from different aphid subfamilies (Chong *et al*., 2019; Gil *et al*., 2006; Van Ham *et al*., 2000). In addition, similar proteins have been found in the pBioThi plasmids of the co-obligate symbiont *Er. haradaeae*, which show a duplicated inverted region causing a duplication of biotin- and thiamin-biosynthetic genes (Manzano-Marín *et al*., 2020). Therefore, these DNA-resolvase/invertases could play a role in rearrangement of plasmidic genes in different endosymbiotic lineages. Both of these novel genes were likely acquired through horizontal gene transfer (HGT) events, from a hitherto unknown origin. HGT events have been observed in the aphid endosymbiont *Er. haradaeae*, where biotin- and thiamin-biosynthetic genes have been likely horizontally transferred from a once co-existing symbiont related to *Sodalis* bacteria (Manzano-Marín *et al*., 2020). Similar cases of HGT have been observed in other small vertically transmitted symbiont genomes. These include the aforementioned example of *Er. haradaeae*, the transfer of an antifungal lagriamide biosynthetic gene cluster in the *Burkholderia* symbiont of the beetle *Lagria villosa* (Waterworth *et al*., 2020), and the acquisition of a biotin-biosynthetic gene cluster in *Wolbachia* (Driscoll *et al*., 2020; Gerth and Bleidorn, 2017; Nikoh *et al*., 2014), *Legionella polyplacis*, (Říhová *et al*., 2017), and the *Midichloria* symbiont of the tick *Hyalomma marginatum* (Buysse *et al*., 2021), among others. HGT events in *Buchnera* and other endosymbionts evidence that, despite a degraded machinery needed for homologous recombination and their highly compact and degraded genomes, these symbionts are still able to horizontally acquire genes from unrelated bacteria. While the route(s) facilitating this HGT events remain, to our knowledge, unknown, the co-infection of bacteriocyte cells by facultative bacteria (Fukatsu *et al*., 2000) could represent an opportunity for such an event.

In conclusion, we found that co-obligate symbiotic systems are more widespread in aphids than previously thought, potentially existing in at least 11.4% of aphid species, given known species diversity in aphid lineages that are now inferred as hosting a dual nutritional symbiosis. One recent study corroborates our finding of a co-obligate *Se. symbiotica* endosymbiont in *Si. maydis* (Renoz *et al*., 2022b), and another one shows the evolution of a co-obligate symbiotic system in *Ceratovacuna japonica* (Hormaphidinae: Cerataphidini; Yorimoto *et al*., 2022). These results, in light of the large-scale approach taken in this work, show that these multi-partner symbioses have evolved several times independently, even within subfamilies. A few major aphid lineages are still missing from this reconstruction, most markedly members of two well-diversified subfamily Greenideinae and Hormaphidinae where more dual symbiotic systems might be revealed. Our analyses also show that a few key gene losses (i.e. those in the riboflavin biosynthetic pathway), rather than large-scale ones, are at the onset of co-obligate symbioses and that genome reduction can be decoupled from new nutritional symbiont acquisitions. It has sometimes been suggested that it is the pervasiveness of facultative symbionts the one that relax selective pressure on genes from *Buchnera* and make it plunge further down the rabbit hole of endosymbiosis leading to co-dependency on a third partner (Lamelas *et al*., 2011a). Our data on *Buchnera* genome degradation, co-obligate symbiont acquisition, and symbiont prevalence across subfamilies, gives support to a scenario where it is not the presence of a facultative symbiont that triggers nor accelerates *Buchnera* decay by relaxing selective constraints of this obligate symbiont. Given the ongoing *Buchnera* genome degradation, even in mono-symbiotic systems (reported here and in Chong *et al*., 2019), it is likely that the decay of *Buchnera* genomes is inherent to its lifestyle, and when it affects gene loss and enzymatic activity, it opens a niche for the reliance on a new symbiont: likely any bacterial lineage that is present and potentially capable of filling that niche. We anticipate that analyses using a larger dataset (*i.e.* including more aphid subfamilies and deeper genomic investigations into each clade) will allow to formally test if *Buchnera* genome characteristics change significantly upon the acquisition of a co-symbiont. This could be achieved by, for example, testing whether substitution models vary in branches of the *Buchnera* tree where dual-symbiosis occurs.

Finally, the role of additional or lost metabolic capabilities (such as the reacquisition of the thiamin biosynthetic pathway) in aphid symbiotic systems remains to be explored. We expect that comparative analyses into ingested phloem by specific aphid species as well as assessment of the specific dependence on nutrients by aphids at different life stages will shed light on whether these gain/losses of metabolic potential can facilitate the establishment of new obligate symbionts.

## Materials and Methods

### Aphid collection and sequencing

Twenty-five different species of aphids were sourced from the *CBGP - Continental Arthropod Collection* (Centre de Biologie pour la Gestion des Populations, 2018; supplementary table S3, Supplementary Material online), where the specimens were preserved in ethanol 70% at 6°C until extraction. In this collection, a specimen (with its unique voucher) corresponds to individuals from a single aphid colony. Bacteria-enriched DNA extractions were performed following Jousselin *et al*. (2016). When possible, 10-15 individuals were used for extraction.

For genomic sequencing, extracted DNA was used to prepare two custom paired-end libraries in Genoscope. Briefly, 5ng of genomic DNA were sonicated using the E220 Covaris instrument (Covaris, USA). Fragments were end-repaired, 3’-adenylated, and NEXTflex PCR free barcodes adapters (Bioo Scientific, USA) were added by using NEBNext Ultra II DNA library prep kit for Illumina (New England Biolabs, USA). Ligation products were purified by Ampure XP (Beckman Coulter, USA) and DNA fragments (*>*200bp) were PCR-amplified (2 PCR reactions, 12 cycles) using Illumina adapter-specific primers and NEBNext Ultra II Q5 Master Mix (NEB). After library profile analysis by Agilent 2100 Bioanalyser (Agilent Technologies, USA) and qPCR quantification using the KAPA Library Quantification Kit for Illumina Libraries (Kapa Biosystems, USA), the libraries were sequenced using 251 bp paired-end reads chemistry on a HiSeq2500 Illumina sequencer. In addition, Nanopore sequencing was done for the aphid species *Anoecia corni*. For long-read sequencing, the Rapid Low Input by PCR Sequencing Kit (SQK RLI001) was used. The library was sequenced on an R9.4 flow cell. In addition to the newly produced sequencing reads, data for an additional 19 species was downloaded from the NCBI’s Short Read Archive (supplementary table S4, Supplementary Material online), with a selection based on two criteria: sequencing data quality and phylogenetic coverage of aphid subfamilies. We therefore did not include all of the previously reported symbiont genomes of Lachninae, as the co-obligate associations have been extensively investigated in previous studies, nor Aphidinae, which so far have only been shown to host *Buchnera* as an obligate nutritional symbiont.

For amplicon sequencing, we investigated symbiont diversity in 223 aphid samples, comprised of 147 species of 75 genera belonging to 12 subfamilies sourced from the aforementioned Aphididae collection. These species included those used for symbiont genome data as well as species closely related to them. DNA was extracted from single individuals and a 251bp portion of the V4 region of the 16S rRNA gene was chose for amplification following Mizrahi-Man *et al*. (2013). The 16S fragment was amplified using primers 16S-V4F (5’-GTGCCAGCMGCCGCGGTAA-3’) and 16S-V4R (5’-GGACTACHVGGGTWTCTAAT-3’) following the dual-index sequencing strategy developed by Kozich *et al*. (2013) and the protocol described in Jousselin *et al*. (2016). Each DNA extract was amplified twice along with negative controls (DNA extraction and PCR controls) using distinct 96-well microplates for PCR replicates. We obtained a total of 485 PCR products, which were pooled together and subjected to gel electrophoresis. The bands corresponding to the PCR products were excised from the gel, purified with a PCR clean-up and gel extraction kit (Macherey-Nagel) and quantified with the Kapa Library Quantification Kit (Kapa Biosystems). The DNA pool was then paired-end sequenced on an Illumina MiSeq flowcell with a 500-cycle Reagent Kit v2 (Illumina).

### 16S amplicon sequencing and analysis

16S amplicon sequences were first filtered through Illumina’s quality control procedure. We then used a pre-processing script from Sow *et al*. (2019) which uses FLASH v1.2.11 (Magoč and Salzberg, 2011) and CUTADAPT v1.9.1 (Martin, 2011) to merge paired sequences into contigs and trim primers, respectively. We then used the FROGS pipeline (Escudié *et al*., 2018) to generate an abundance table of symbiont lineages across samples. Briefly, to generate the tables, we first filtered out sequences >261 and <241 bp. We then clustered variants with Swarm (Mahé *et al*., 2014) using a maximum aggregation distance of 1. Lastly, we identified and removed chimeric variants with VSEARCH (Rognes *et al*., 2016).

Taxonomic assignments of clusters was carried out using RDPtools v2.0.3 (https://github.com/rdpstaff-/RDPTools, last accessed July 18, 2022; Cole *et al*., 2014) and BLASTn+ (Camacho *et al*., 2009) against the Silva database release 138 (Quast *et al*., 2013) as implemented in FROGS. Following taxonomic affiliation, we aggregated clusters when they shared the same taxonomy with at least 98% of identity (FROGS’ affiliation postprocess step). From the abundance table of clusters across samples, we transformed read numbers per aphid samples into percentages and sequences accounting for *<*0.5% of all the reads for a given sample were excluded using an R script following Jousselin *et al*. (2016). Clusters were kept only if present in both PCR replicates of the same sample. For final description of endosymbiont diversity we only kept the PCR replicate that yielded the highest number of reads. We refined ambiguous taxonomic assignations using BLASTn+ against 16S rRNA gene sequences extracted from whole endosymbiont genomes from this study and retrieved from GenBank. We also used the webtool leBIBI IV (https://umr5558-proka.univ-lyon1.fr/lebibi/lebibi.cgi, last accessed July 18, 2022) that provides automatic phylogenetic placement of bacterial 16S sequences (Flandrois *et al*., 2015). The resulting relative abundance table (supplementary table S5, Supplementary Material online) was used to produced a heatmap in R of presence/absence of endosymbiotic taxa across aphid subfamilies. Presence/absence was codified as 0 and 1, respectively. These 0/1 numbers were then scaled to the total number of samples per subfamily to provide a more accurate visual representation of secondary symbiont abundance across the samples in figure 4.

### Genome assembly

For all Illumina datasets, reads were right-tail clipped (using a minimum quality threshold of 20) using FASTX-Toolkit v0.0.14 (http://hannonlab.cshl.edu/fastxtoolkit/, last accessed July 18, 2022) and those shorter than 75 after were dropped. PRINSEQ v0.20.4 (Schmieder and Edwards, 2011) was used to remove reads containing undefined nucleotides as well as those left without a pair after the filtering and clipping process. Clean reads were assembled using SPAdes v3.11.1 (Bankevich *et al*., 2012) with the --only-assembler option and k-mer sizes of 33, 55, 77, 99, and 127 (depending on the read length). Assembled contigs shorter than 200 bps were dropped. The surviving contigs were binned using results from a BLASTx (Altschul *et al*., 1997) search (best hit per contig) against a database consisting of the proteomes of the Pea aphid and a selection of aphid’s symbiotic and expected free-living bacteria (supplementary table S6, Supplementary Material online). When no genome was available for a certain lineage, closely related bacteria were used.

The binned contigs were manually screened using the BLASTx web server (*vs.* the nr database) to insure correct taxonomic assignment. The resulting contigs were then used as reference for read-mapping and individual genome re-assembly using SPAdes, as described above. Binned and re-assembled contigs were checked for circularisation and completion through assembly graphs. For those *Buchnera* strains that have evolved a low-complexity replication start, we closed the ends with ”N” stretches after checking for the conserved proteins around the end and truncation of low-complexity sequencing artefacts. Details of any sequence modification post-assembly is captured in the GenBank-formatted annotation files.

### Genome annotation

The resulting *Buchnera* genomes were annotated as follows. First, open reading frame (ORF) prediction was done using Prokka v1.14.6 (Seemann, 2014). This ORF prediction was followed by non-coding RNA prediction with infernal v1.1.2 (Nawrocki and Eddy, 2013) (against the Rfam v14.1 database; Kalvari *et al*., 2021), tRNAscan-SE v2.0.9 (Chan *et al*., 2021), and ARAGORN v1.2.36 (Laslett, 2004). This annotation was followed by careful manual curation of the genes on UGENE v37.1 (Okonechnikov *et al*., 2012) through on-line BLASTx searches of the intergenic regions as well as through BLASTp and DELTA-BLAST (Boratyn *et al*., 2012) searches of the predicted ORFs against NCBI’s nr database. The resulting coding sequences (CDSs) were considered to be putatively functional if all essential domains for the function were found or if a literature search supported the truncated version of the protein as functional in a related organism, or if the the CDS displayed truncations but retained identifiable domains (details of the literature captured in the annotation file). Pseudogenes were further searched for based on synteny with closely related *Buchnera* strains on which previous manual curation was done. This prediction was performed based on nucleotide sequence using a combination of sequence alignment (with m-coffee; Wallace et al. 2006) and BLASTx searches against the NCBI’s nr database. This last check allowed for the identification of pseudogenes missed by the previous searches. Once a *Buchnera* was manually curated, it was used as a Prokka reference protein set for the next *Buchnera* in order to maintain naming congruent across genomes. Following experimental evidence presented in Tamas *et al*. (2008), genes interrupted by a frameshift in a low complexity A- or T-homopolymeric region, were annotated as coding for a functional protein, with special attention to conservation of frameshifted region and amino acid sequence within closely related *Buchnera* strains.

For the genomes of co-obligate symbionts, draft Prokka annotations were performed and genes of interest to metabolic complementarity for nutrient provisioning underwent a manual curation as described above. For genome statistics, an estimated count for pseudogene features was done as follows. Contiguous proteins detected as truncated and with same assignment as well as proteins under 100 amino acids, were considered pseudogenes. tRNAs, ncRNAs, and rRNAs were predicted as described above.

### Phylogenetics

In order to reconstruct a phylogenetic hypothesis for *Buchnera*, we used OrthoMCL v2.0.9 (Chen *et al*., 2007; Li, 2003) to build clusters of orthologous proteins. To the newly acquired, reassembled, and re-annotated symbiont genomes; we added three already annotated ones from Lachninae (supplementary table S4, Supplementary Material online). These orthologous groups were then manually curated using the annotation to keep same genes together in clusters. Count v10.04 (Csuos, 2010)was used to calculate the most parsimonious scenario for gene losses in *Buchnera*. *Escherichia coli* K-12 MG 1655 and a selection of *Pantoea* and *Eriwnia* strains were used as outgroups (supplementary table S7, Supplementary Material online). We then retrieved the single copy-core proteins of the selected genomes for phylogenetic reconstruction. We aligned the single-copy core protein sets, gene by gene, using MAFFT v7.453 (Katoh and Standley, 2013) (L-INS-i algorithm). Divergent and ambiguously aligned blocks were removed using Gblocks v0.91b (Talavera and Castresana, 2007). The resulting alignments were concatenated for phylogenetic inference. Bayesian inference was performed in MrBayes v3.2.7a (Ronquist *et al*., 2012) running two independent analyses with four chains each for up to 300,000 generations and checked for convergence with a burn-in of 25%. JModelTest v2.1.10 (Darriba *et al*., 2012) was used to select the best model for phylogenetic reconstruction based on the Akaike’s Information Criterion (cpREV+I+G4).

To analyse the phylogenetic relations of newly sequenced aphid co-obligate endosymbionts, we built two phylogenies. We first reconstructed a phylogeny for *Yersinia*- and *Serratia*-related endosymbionts. For this, we used the dataset built by (Roüıl *et al*., 2020) and added data for *Serratia symbiotica* symbionts and the newly sequenced symbiotic strains. Given that many *Se. symbiotica* genomes lacked an intact *hrpA*, this gene was excluded from our dataset. Each gene set was individually aligned using MUSCLE v3.8.31 (Edgar, 2004). Then, we removed divergent and ambiguously aligned blocks with Gblocks. Bayesian inference was performed in MrBayes, using the GTR+I+G4 substitution model running two independent analyses with four chains each for 1,000,000 generations and checked for convergence. The second phylogeny was reconstructed for *Sodalis*-like endosymbionts. Given that many of these endosymbionts remain without a sequenced genome, we used 16S rRNA gene sequences following (Manzano-Marín *et al*., 2017). We used SSU-ALIGN v0.1 (Nawrocki, 2009) to align the rRNA gene sequences. GBlocks v0.91b was then used to eliminate poorly aligned positions and divergent regions with the option ‘-b5=h’ to allow half of the positions with a gap. MrBayes v3.2.7 was used for phylogenetic inference under the GTR+I+G4 model running two independent analyses with four chains each for up to 10,000,000 generations and checked for convergence, discarding the first 25% as burn-in. Visualization and tree-editing for all analyses was done in FigTree v1.4.1 (http://tree.bio.ed.ac.uk/software/figtree/, last accessed July 18, 2022).

### Fluoresence *in situ* hybridisation microscopy

Live aphid specimens from *Sipha maydis* and *Anoecia corni* were fixed in modified Carnoy’s fixative (6 chloroform: 3 absolute ethanol: 1 glacial acetic acid) and left overnight, following the protocol of (Koga *et al*., 2009). Individuals were then dissected in absolute ethanol to extract embryos and transferred into a 6% solution of H_2_O_2_ diluted in absolute ethanol and were then left in this solution for two weeks (changing the solution every three days). Embryos were then washed twice with absolute ethanol. Hybridization was performed overnight at 28°C in standard hybridisation buffer (20 mM Tris-HCl [pH 8.0], 0.9M NaCl, 0.01% SDS, and 30% formamide) and then washed (20 mM Tris-HCl [pH 8.0], 5 mM EDTA, 0.1M NaCl, and 0.01% SDS) before slide preparation. Competitive probes were adapted for this specific symbionts from (Manzano-Marín *et al*., 2017) (supplementary table S8, Supplementary Material online).

All files relating to orthologous protein grouping as well as phylogenetic reconstructions can be found at https://doi.org/10.5281/zenodo.6394197.

## Supplementary Material & Data Availability

Supplementary figures S1-S3 and tables S1-8 and have been included in this submission. All auxiliary files for phylogenetics and other analyses, unmerged FISH microscopy images, as well as the genomes of *Buchnera* and co-obligate endosymbionts are available online at https://doi.org/10.5281/zenodo.6394197. Newly sequenced and annotated genomes are in the process of being submitted for accessioning to the European Nucleotide Archive (ENA).

## Acknowledgements

We would like to acknowledge the talented artist/scientist Jorge Mariano Collantes Alegre for the aphid cartoons in figure 7. We would also like to thank Prof. Aharon Oren for his help and advice in bacterial species nomenclature. This work was supported by the Marie-Curie AgreenSkills+ fellowship programme co-funded by the EU’s Seventh Framework Programme (FP7-609398) to A.M.M. The Marie-Skłodowska-Curie H2020 Programme (H2020-MSCA-IF-2016) to E.J., the Agropolis foundation/Labex Agro (”Cinara’s microbiome”) to E.J, the France Génomique National Infrastructure, funded as part of the *Investissement d’Avenir* program managed by the *Agence Nationale pour la Recherche* (ANR-10-INBS-0009) to E.J., C.C., and V.B. This publication has been written with the support of the AgreenSkills + fellowship programme which has received funding from the EU’s Seventh Framework Programme under grant agreement No. FP7-609398 (AgreenSkills + contract) and the Horizon 2020 Programme under grant agreement 746189 (H2020-MSCA-IF-2016-MicroPhan). We are grateful to the genotoul bioinformatics platform Toulouse Midi-Pyrenees (Bioinfo Genotoul) for providing help and/or computing and/or storage resources. The authors are grateful to the CBGP-HPC computational platform and the staff maintaining the *Life Science Compute Cluster* (LiSC; http://cube.univie.ac.at/lisc) at the University of Vienna which were used for computational analysis. The funders had no role in study design, data collection and analysis, decision to publish, or preparation of the manuscript.

