## Supplementary figures S1-S4 for "Co-obligate symbioses have repeatedly evolved across aphids, but partner identity and nutritional contributions vary across lineages"

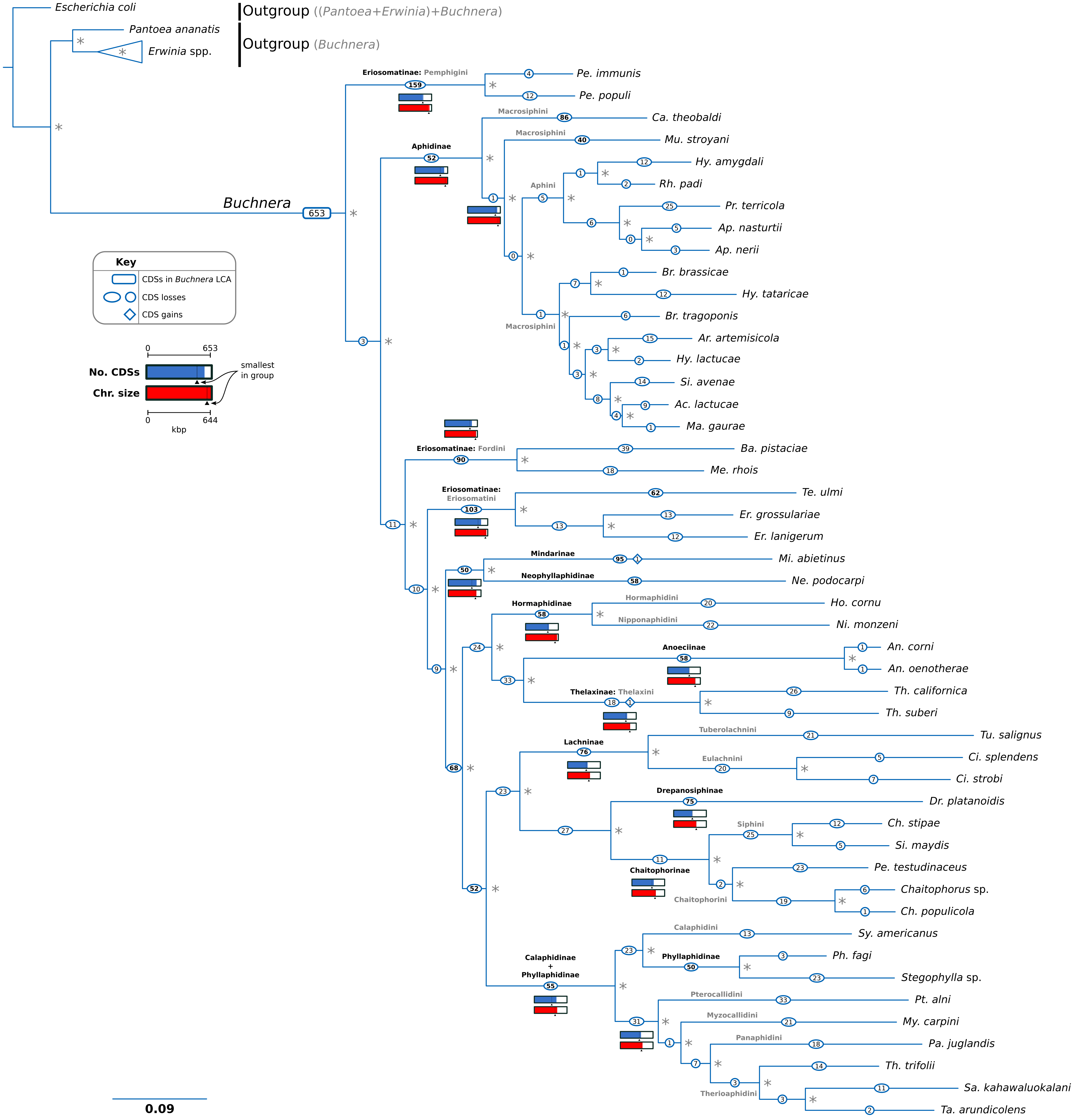


**Figure S1. Phylogenetic relationships of *Buchnera*.** Bayesian phylogenetic tree based on concatenated single-copy core proteins of selected *Buchnera*. *Escherichia coli* K-12 MG1655 and selected *Pantoea* and *Erwinia* strains were used as outgroups to root the tree. Names of aphid subfamilies and tribes are shown in bold font in black and grey, respectively. Bar representations at selected nodes display number of CDSs and genome sizes of the *Buchnera* genomes in that phylogenetic group. Given that *fliM+fliN* and *fliO+fliP* often occurred as fused genes, they were each counted as one protein cluster instead of two. An asterisk at nodes stands for a posterior probability of 1.

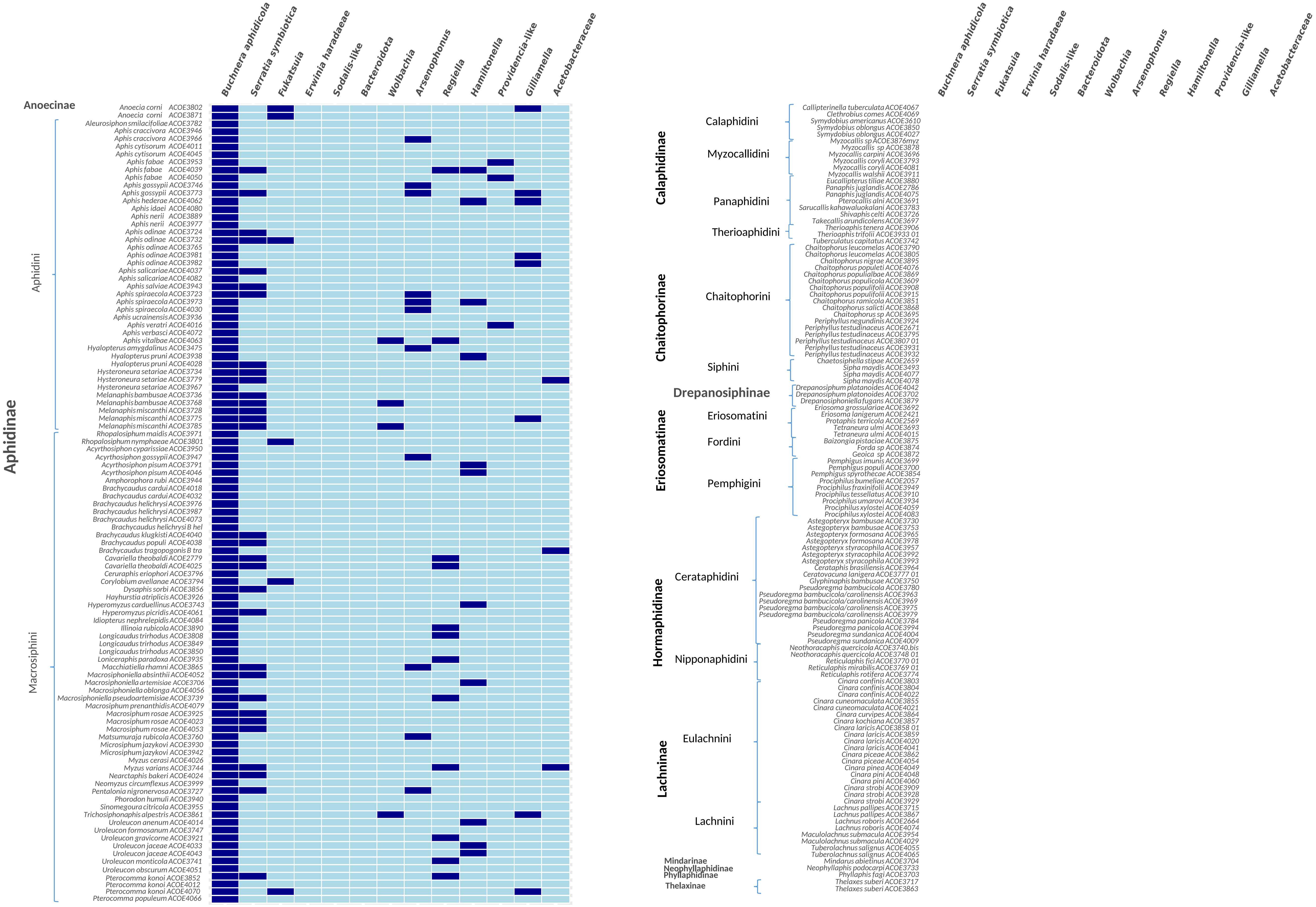


**Figure S2. Presence/absence of bacterial taxa across aphid taxa.** Matrix summarising results of 16S rRNA amplicon taxonomic assignment of reads in the form of presence (dark blue)/absence (light blue). Each row and column represents an aphid sample and a bacterial taxon, respectively. On the left, subfamily- and tribal-level groupings are shown.

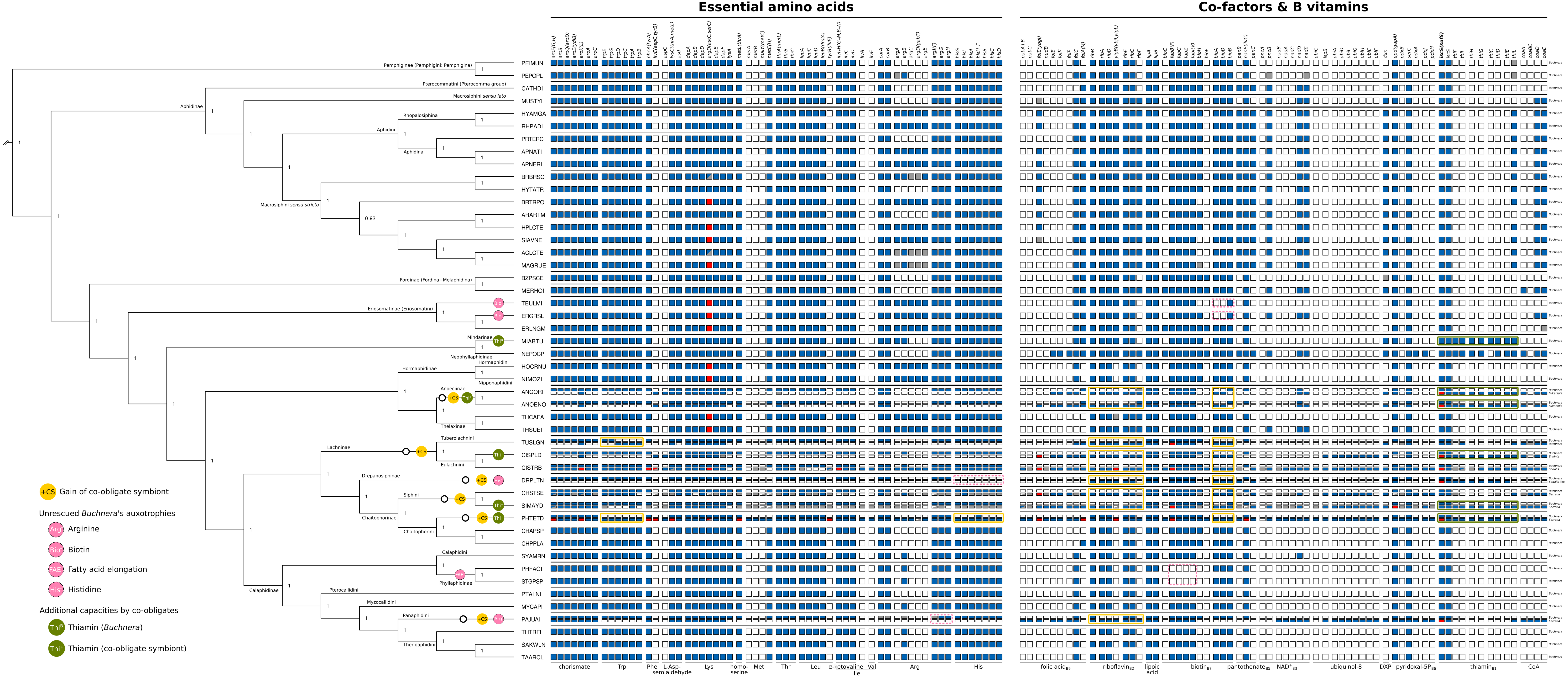


**Figure S3. Detailed metabolic complementarity of co-obligate symbiotic systems of aphids.** Matrix showing metabolic capacities of obligate symbionts of different aphid species. On the left, dendrogram displaying phylogenetic relationships of *Buchnera* strains (see supplementary figure S1). An empty circle at a node represent the acquisition of a co-obligate symbiont. At the leaves, abbreviation for aphid taxa. At the top of each box column, the name of the enzymes or pathway catalysing a reaction. At the bottom, the name of the compound synthesised by the enzymatic steps. At the right of each row of boxes, symbiont taxon name. Black and grey bars between boxes separate aphid subfamilies and tribes. PEIMUN= *Pe. immunis*, PEPOPL= *Pe. populi*, CATHDI= *Ca. theobaldi*, MUSTYI= *Mu. stroyani*, HYAMGA= *Hy. amygdali*, RHPADI= *Rh. padi*, PRTERC= *Pr. terricola*, APNATI= *Ap. nasturtii*, APNERI= *Ap. nerii*, BRBRSC= *Br. brassicae*, HYTATR= *Hy. tataricae*, BRTRPO= *Br. tragopogonis*, ARARTM= *Ar. artemisicola*, HPLCTE= *Hy. lactucae*, SIAVNE= *Si. avenae*, ACLCTE= *Ac. lactucae*, MAGRUE= *Ma. gaurae*, BZPSCE= *Ba. pistaciae*, MERHOI= *Me. rhois*, TEULMI= *Tetraneura ulmi*, ERGRSL= *Er. grossulariae*, ERLNGM= *Er. lanigerum*, MIABTU= *Mi. abietinus*, NEPOCP= *Ne. podocarpi*, HOCRNU= *Ho. cornu*, NIMOZI= *Ni. monzeni*, ANCORI= *An. corni*, ANOENO= *An. oenotherae*, THCAFA= *Th. californica*, THSUEI= *Th. suberi*, TUSLGN= *Tu. salignus*, CISPLD= *Ci. splendens*, CISTRB= *Ci. strobi*, DRPLTN= *Dr. platanoidis*, CHSTSE= *Ch. stipae*, SIMAYD= *Si. maydis*, PHTETD= *Pe. testudinaceus*, CHAPSP= *Chaitophorus* sp., CHPPLA= *Ch. populicola*, SYAMRN= *Sy. americanus*, PHFAGI= *Ph. fagi*, STGPSP= *Stegophylla* sp., PTALNI= *Pt. alni*, MYCAPI= *My. carpini*, PAJUAI= *Pa. juglandis*, THTRFI= *Th. trifolii*, SAKWLN= *Sa. kahawaluokalani*, TAARCL= *Ta. arundicolens*. At nodes, values for posterior probabilities is shown.

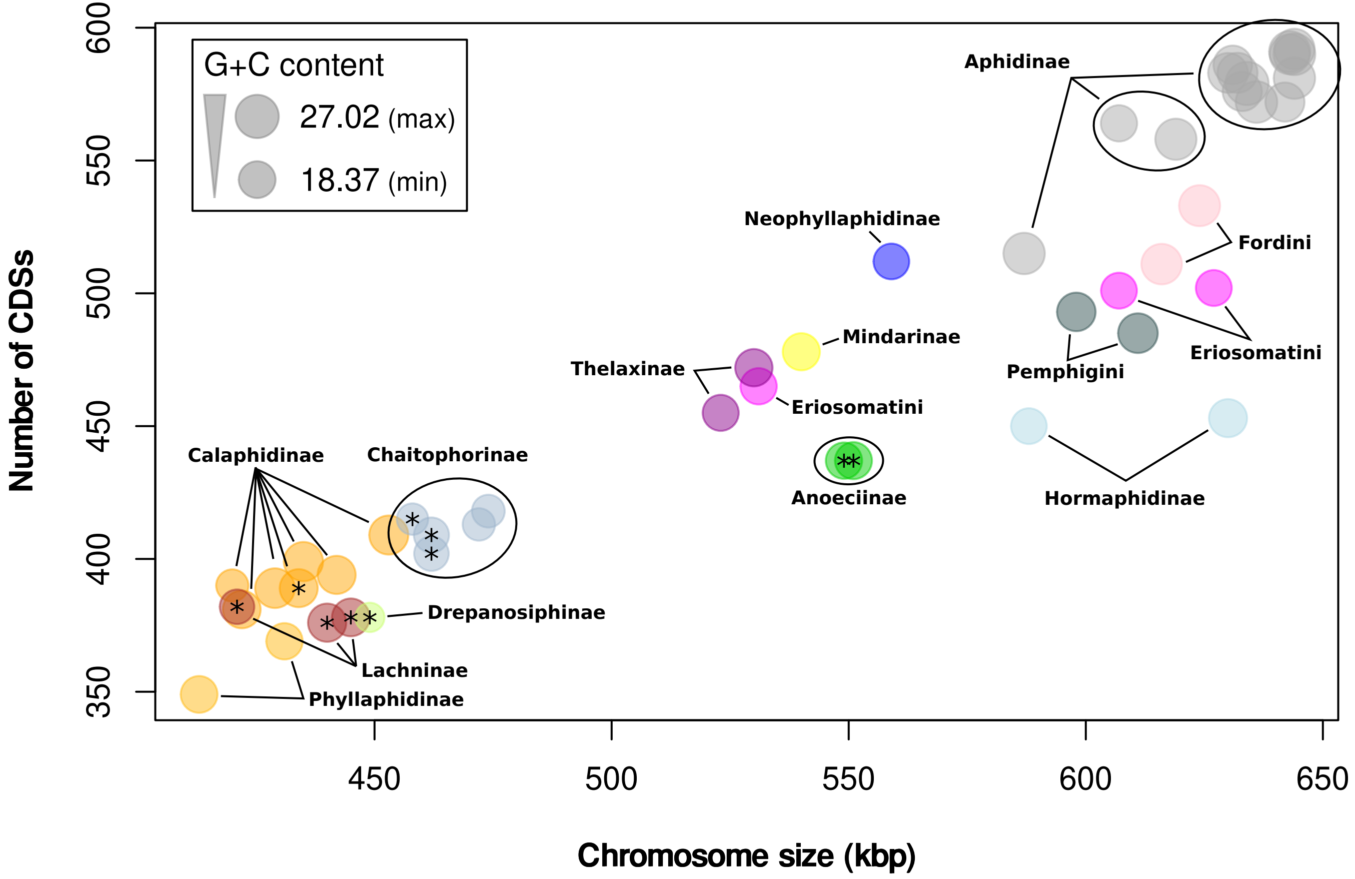


**Figure S4. Relationship of genome size *vs.* CDS content in *Buchnera* genomes.** Scatter plot illustrating the diversity of genome size, number of CDSs and G+C content of selected *Buchnera* genomes. Taxonomic classification of aphid host is displayed in bold black lettering. Dots are colour coded according to the host's subfamily or tribal classification. Size of circles represent the G+C content as shown in the legend box scale key. An "*" highlights *Buchnera* strains sharing their aphid host with a co-obligate endosymbiont.
